# Extracellular Vesicle-Mediated Delivery of HIF1α Reprograms Macrophages for resolutive response in Sepsis

**DOI:** 10.1101/2024.08.29.608900

**Authors:** Yeji Lee, Jiyoung Goo, Seongeon Cho, Seong A Kim, Gi-Hoon Nam, Iljin Kim, Jong-Wan Park, Cherlhyun Jeong, In-San Kim

**Affiliations:** KU-KIST Graduate School of Converging Science and Technology, Korea University, Seoul 02841, Republic of Korea; KHU-KIST Department of Converging Science and Technology, Kyunghee University, Seoul 02447, Republic of Korea; Chemical and Biological Integrative Research Center, Korea Institute of Science and Technology, Seoul 02792, Republic of Korea; Division of Bio-Medical Science & Technology, University of Science and Technology (UST), Seoul 02792, Republic of Korea; SHIFTBIO INC., Seoul, Republic of Korea; Department of Biochemistry and Molecular Biology, Korea University College of Medicine, Seoul, Republic of Korea; Department of Pharmacology and Program in Biomedical Science and Engineering, Inha University College of Medicine, Incheon 22212, Republic of Korea; Department of Pharmacology, Seoul National University College of Medicine, Seoul 03080, Republic of Korea

## Abstract

In sepsis, the liver functions as a central filter organ, where hepatic macrophages form a primary antimicrobial defense layer by eliminating bacteria and regulating immune responses. Therefore, precise regulation of the immune response in hepatic macrophages is crucial for triggering effective defense mechanisms. We aim to modulate the defense immune response by delivering transcription factor HIF1α, a key regulator of monocyte/macrophage reprogramming in sepsis. Transcription factors are promising candidates because they dynamically modulate gene expression across diverse conditions, though delivering them remains challenging. In this study, we suggest a novel method for loading HIF1α into extracellular vesicles (EVs) to enhance immune defense and resolve sepsis. By delivering HIF1α to macrophages during sepsis, we promoted the differentiation of Nr1h3-dependent pro-efferocytic MoMFs and C/ebpβ-dependent pro-survival MoMFs. Pro-efferocytic MoMFs eliminate damaged hepatocytes and immune cells and pro-survival MoMFs withstand inflammatory conditions and trigger innate memory responses. Particularly, the zonation of these macrophages in the periportal region ensures effective pathogen clearance and minimizes tissue damage. These findings suggest that EV-mediated HIF1α delivery could be a promising therapeutic option for managing sepsis.

## main

Sepsis is a complex complication arising from infections caused by bacteria, viruses, and parasites, leading to excessive immune responses and systemic inflammation, resulting in multi-organ dysfunction.^1–5^ Particularly, the liver is a critical filter organ responsible for about 60% of endotoxin and bacteria clearance.^6,7^ Therefore, liver dysfunction is considered an independent predictor of mortality in sepsis.^8–10^ Patients with liver injury or cirrhosis have compromised defense mechanisms against microbial infections, making effective bacterial clearance difficult.^2,4,7,8^ This can lead to excessive inflammation or induce chronic inflammation in the liver.

The liver has a high density of innate immune cells, enabling rapid activation of immune responses in reaction to tissue damage or infection.^3,10,11^ In early infection, resident immune cells like Kupffer cells, LSECs, and HSCs induce a fast response by recruiting neutrophils and macrophages.^12,13^ The recruited immune cells initiate immune responses to eliminate the pathogens, but this can also perpetuate inflammation and cause additional tissue damage.^8,12^ Consequently, reprogramming the recruited immune cells to induce an efficient immune response and resolve inflammation is crucial. However, in the early stages where inflammation is excessively activated, heterogeneous immune cells infiltrate the liver, making it challenging to regulate and reprogram them. Nevertheless, signaling pathways in complex environments generally converge on a single transcription factor (TF) in a bowtie topology.^14^ Therefore, we believe that a single transcription factor could precisely regulate the complex immune response and induce a resolutive environment.

HIF1α is a transcription factor regulated by oxygen levels, being degraded under normal oxygen conditions and accumulated in hypoxic and inflammatory environments.^15–17^ This suggests that HIF1α needs to respond immediately and rapidly under hypoxic and inflammatory conditions, even at the cost of maintaining an inefficient biosynthetic process.^17^ Particularly, HIF1α has been identified as a key mediator in reprogramming monocytes and macrophages during sepsis.^16,18^ HIF1α in monocytes/macrophages has been shown to upregulate genes related to antimicrobial activity and tissue remodeling.^16^ Additionally, in the early stages of liver injury, HIF1α expression in monocyte-derived macrophages (MoMFs) promotes necrotic tissue removal and induces liver repair.^18^ In this study, we hypothesized that delivering HIF1α could reprogram monocytes/macrophages, which play a central role in sepsis, to induce a resolutive environment.

Despite these advantages, the main challenge remains in efficiently transporting TFs across the plasma membrane and into the nucleus. Strategies using CPPs and nanovesicles have been developed to facilitate intracellular protein delivery, but they struggle with endosomal escape and toxicity.^19,20^ To overcome these challenges, we aimed to deliver the TF using extracellular vesicles (EVs), natural delivery vehicles known for their lack of hepatotoxicity and efficient endosomal escape.^21–23^

## Result

### Efficient Delivery of HIF1α via EVs Using the Self-Cleaving Platform

First, we propose new strategy to deliver the transcription factor (TF) HIF1α using Extracellular Vesicles (EVs) (Fig 1A). To effectively deliver HIF1α, it is essential to comprehend its intrinsic properties of TFs and HIF1α. Once expressed in the cytoplasm, TFs generally translocate to the nucleus. Therefore, in the parent cells, the TF should be sorted into the EVs without entering the nucleus, and in the recipient cells, the TF should exist in a free form to facilitate nuclear translocation. We connected the pH-dependent Self-Cleaving intein System (I-CM)^24^ between the CD81 which is a EV marker and HIF1α, allowing HIF1α to undergo cleavage after delivery into the EVs and exist in a free form within the low pH environment of the EVs.^25^ HIF1α stabilizes under hypoxic conditions but rapidly degrades in normoxic conditions, making it difficult to maintain its structural stability during EV production. Therefore, we created a stable construct of HIF1α (scHIF1α) by removing the ODD domain^26^, which binds hydroxyl groups in response to oxygen and promotes degradation by protease, allowing it to remain stable even in normoxic conditions (Fig 1B).

**Fig. 1.**
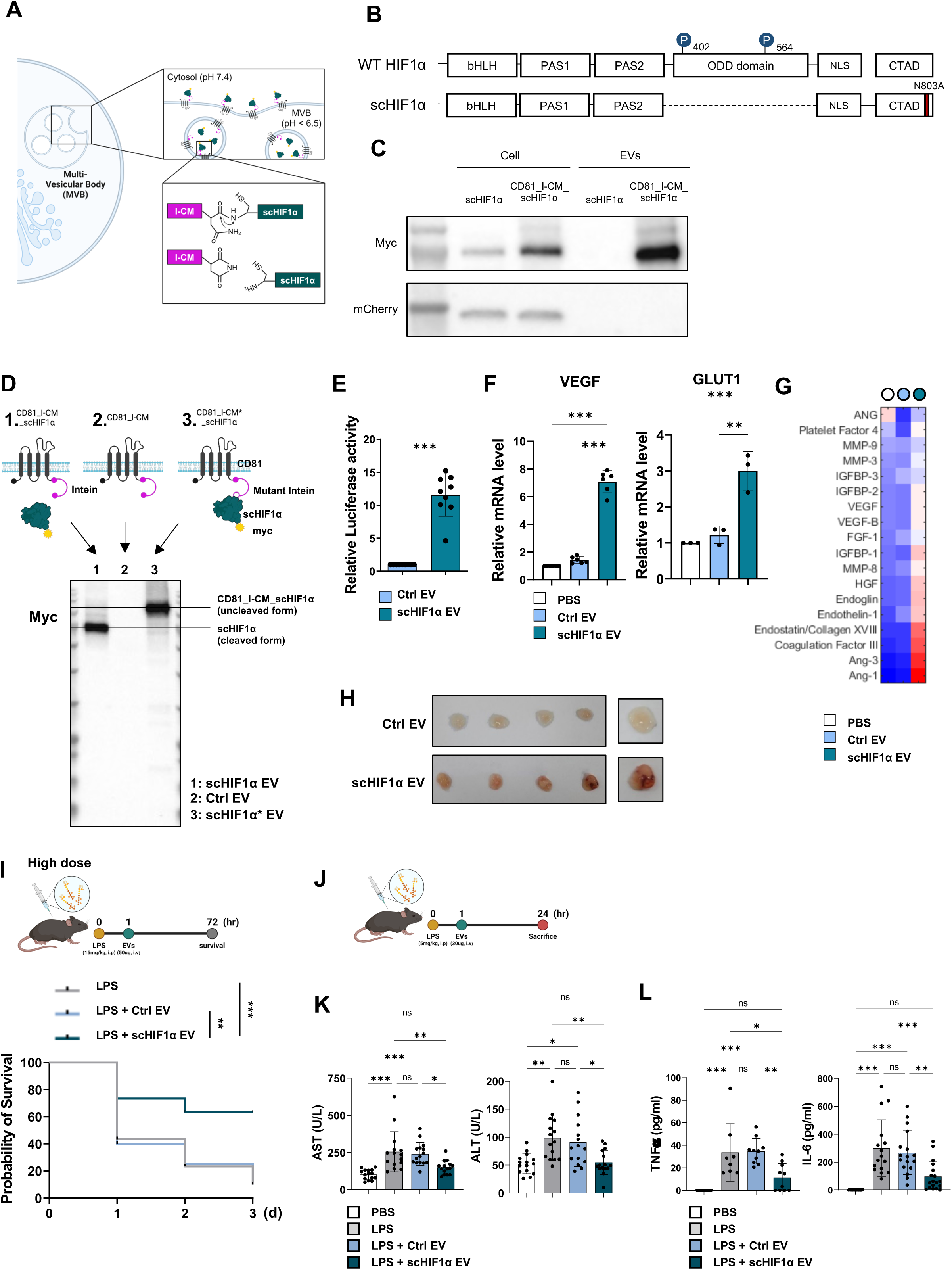
Overview of the strategy for delivering the transcription factor HIF1α and the verification of nuclear delivery effects and in vivo efficacy. A. Illustration of the HIF1α delivery mechanism using pH-dependent mini intein. B. Structures of wild-type HIF1α (top) and ODD domain-deleted scHIF1α (bottom). C. Validation of scHIF1α loading efficiency via WB. HEK293T cells were transfected with scHIF1α (plasmid without the CD81_I-CM platform) and CD81_I-CM_scHIF1α. mCherry was co-transfected to confirm transfection efficiency. D. Verification of the cleavage effect of pH-dependent mini intein by Western blot. EVs obtained from transfected structures; CD81_I-CM_scHIF1α (Lane 1), CD81_I-CM (Lane 2), CD81_I-CM*_scHIF1α (Lane 3). E. Verification of nuclear delivery of scHIF1α by measuring relative luciferase activity. Luciferase activity was assessed in HEK293T cells transfected with the VEGF-luc plasmid and treated with scHIF1a EVs and Ctrl EVs. F. Verification of nuclear delivery of scHIF1α by qRT-PCR of downstream gene expression. Expression of HIF1α downstream genes VEGF and GLUT1 in HEK293T cells treated with scHIF1α EVs and Ctrl EVs was confirmed by qRT-PCR. G. Confirming scHIF1α-mediated angiogenesis via cytokine array. Liver macrophages were sorted using F4/80+ beads and treated with scHIF1α EVs. H. Confirming scHIF1α-mediated angiogenesis via in vivo matrigel assay. A matrigel containing 100 µg of EVs was subcutaneously injected into BALB/c nude mice. I. Confirmation of survival rates by scHIF1α EVs in an LPS-induced sepsis model. LPS (15 mg/kg) was administered via intraperitoneal injection, followed by intravenous injection of 50 µg of scHIF1α EVs. (n=35, 20, 30) J. Experimental design for studying LPS-induced liver damage. LPS (5 mg/kg) was administered by intraperitoneal injection, followed by treatment with scHIF1α EVs one hour later, and analysis was performed after 24 hours. K. Serum ALT and AST levels from mice with LPS-induced sepsis model. (n=15) L. Serum pro-inflammatory cytokine levels, TNFa (n=10) and IL-6 (n=16), from mice with LPS-induced sepsis model. Each column displays group means with individual data points and error bars with SEM. Statistical significance was determined using t-test and one-way ANOVA followed by Tukey’s multiple comparison test (F,K,L). P values indicate significant differences (*p<0.033; **p<0.002; ***p<0.001).

We purified EVs following the MISEV2018 guidelines^27^ and confirmed their size and markers using NTA, cryo-TEM, and Western blot analysis (Extended Fig 1A, B). To verify the sorting efficiency of scHIF1α in EVs, we compared it with the free form of scHIF1α, which is not conjugated to the CD81_I-CM platform. As expected, the scHIF1α was expressed inside the cells but was not sorted into EVs. However, with the CD81_I-CM platform, scHIF1α was predominantly sorted into EVs. To make sure that the difference in scHIF1α expression within cells was not due to transfection efficiency, we co-transfected mCherry, confirming the same transfection efficiency (Fig 1C). We performed a Western blot to identify whether scHIF1α existed in a free form after being cleaved from CD81_I-CM in response to the low pH inside the EVs. To verify cleavage activity within the CD81_I-CM platform, we engineered N168A and C169A mutations to produce a non-cleaving form, CD81_I-CM* (Extended Fig 1C). We compared the molecular weight band of scHIF1α by Western blot of scHIF1α EVs loaded with CD81_I-CM-scHIF1α, Ctrl EVs loaded with CD81_I-CM, and scHIF1α* EVs loaded with CD81_I-CM*-scHIF1α. The non-cleaving construct, CD81_I-CM*_scHIF1α, displayed a band at 145 kDa, indicating no cleavage. In contrast, the cleaving form, CD81_I-CM_scHIF1α, was cleaved inside the EVs, resulting in the detection of the free form of scHIF1α at 75 kDa (Fig 1D). We efficiently sorted the scHIF1α into EVs using the CD81_I-CM platform, and confirmed that it was cleaved inside the EVs, existing in a free form.

To confirm that scHIF1α loaded into EVs is delivered to the nucleus of target cells and functions as a transcription factor, we measured luciferase activity driven by the VEGF promoter (Fig 1E). Additionally, we confirmed the induction of HIF1α downstream genes VEGF and GLUT1 via RT-PCR (Fig 1F) and observed the release of angiogenesis cytokines from media obtained F4/80 sorted macrophages with scHIF1α EVs (Fig 1G). To confirm whether scHIF1α functions as a transcription factor in vivo, we inoculated scHIF1α EV-mixed matrigel subcutaneously into Balb/c nude mice and observed angiogenesis (Fig 1H). To directly confirm the nuclear delivery of TF cleaved by I-CM, we created a TF-mimicking construct 3xNLS_3xsfGFP_2xNLS and observed the delivery to the nucleus in HeLa cells using confocal imaging (Extended Fig 1D). These results demonstrate that scHIF1α loaded into EVs is effectively delivered into the nucleus of recipient cells and functions as a transcription factor in vitro, ex vivo, and in vivo.

### scHIF1α EVs Demonstrate Therapeutic Efficacy in Sepsis-Induced Liver Injury

We evaluated the therapeutic efficacy of scHIF1α EVs in the LPS-induced sepsis model. First, to evaluate the survival rate, we administered a high dose (15 mg/kg) of LPS via intraperitoneal (I.P) injection, followed by an intravenous (I.V) injection of 50 µg of scHIF1α EV one hour later. The survival rate of the scHIF1α EV group showed a significant increase (Fig 1I). Next, we administered a non-lethal dose (5 mg/kg) of LPS to the mice to evaluate liver damage and confirm the immune profile (Fig 1J). The levels of liver damage indicators, AST and ALT^28^, were significantly lower in the scHIF1α group, comparable to the PBS group (Fig 1K). Additionally, a TUNEL assay revealed that the proportion of TUNEL-positive apoptotic cells in the liver tissue was significantly reduced in the scHIF1α group (Extended Fig 2A). The quantification of major inflammatory cytokines^5,23^ TNF-a and IL-6 in serum by ELISA also revealed lower levels in the scHIF1α EV group (Fig 1L). Furthermore, a comprehensive cytokine array conducted in serum indicated reduced cytokine release in the scHIF1α EV group (Extended Fig 2B). These results indicate that scHIF1α EVs can minimize liver injury and improve systemic inflammation in sepsis conditions.

### Systemically Delivered scHIF1α EVs Accumulate Macrophages in Liver

To confirm biodistribution of EVs, we labeled EVs with NHS-Cy5 and administered them via I.V injection, confirming substantial accumulation in the liver (Fig 2A). This was consistent with previous reports showing substantial accumulation of EVs in the liver.^21,22^ To determine which cells in the liver the EVs accumulate in, we used Ai14 reporter mice that express TdTomato upon Cre recombinase uptake and excision of stop cassettes.^29^ We injected EVs loaded with Cre recombinase via I.V and then collected tissues to identify the cells expressing TdTomato. TdTomato expression was predominantly observed in F4/80 positive cells (Fig 2B). To identify the cell type specific biodistribution, we performed FACS analysis after injecting Cy5-labeled EVs via I.V injection. The EVs were predominantly taken up by macrophages and LSECs (Extended Fig 3A). These findings indicate that intravenously injected EVs are dominantly delivered to liver macrophages.^23^ In sepsis-induced liver injury, macrophages are recruited early to secrete inflammatory cytokines to eliminate pathogens,^3^ but this process is also known to cause tissue damage.^8,12^ To evaluate the impact of scHIF1α EVs on cytokine release from macrophages, we injected EVs into a sepsis-induced liver injury model, then isolated macrophages to examine the released cytokines (Extended Fig 3B). Compared to the LPS group, the scHIF1α EVs group showed a relatively increased release of anti-inflammatory cytokines.

**Fig. 2.**
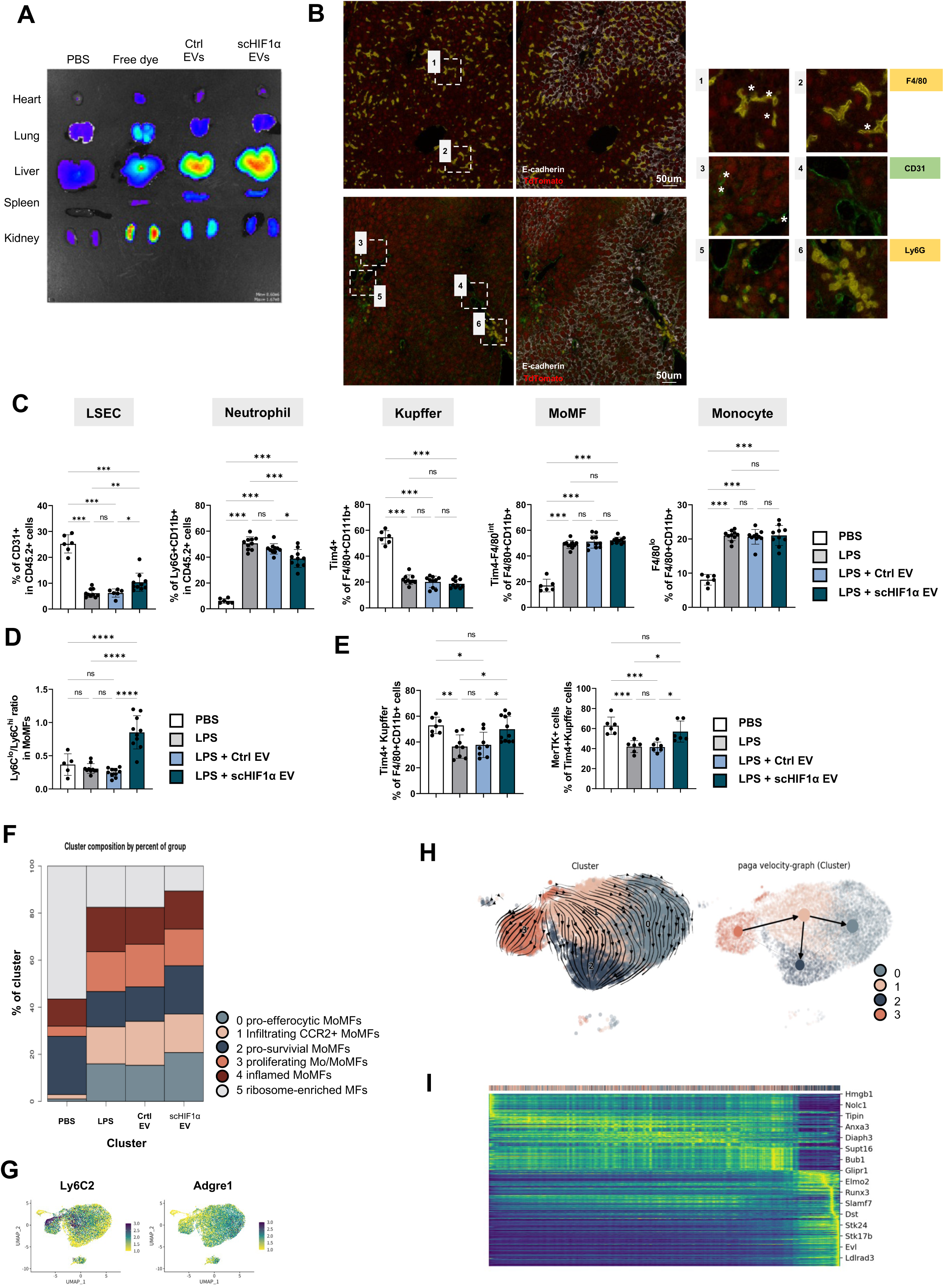
Liver macrophages undergo differentiation upon uptake of scHIF1α EVs. A. Biodistribution imaging after intravenous injection of EVs labeled with Cy5. Images show intensity in the heart, lungs, liver, spleen, and kidneys, from top to bottom. B. Ai14 mice were injected intravenously with 30 µg each of Cre EVs and scHIF1α EVs, and tissues were collected 5 days later for multiple IHC staining. The asterisk (*) indicates TdTomato positive. C. Immune cell profiling by FACS analysis in an LPS-induced sepsis model. The proportion of each immune cell type among total immune cells was analyzed (from left to right: LSEC, Neutrophil, Kupffer (Tim4+), MoMF (Tim4-F4/80int), Monocyte (Tim4-F4/80lo)). (n=6, 10, 10, 10) D. Graph showing the Ly6Clo/hi ratio in MoMFs from FACS analysis. (n=6, 10, 10, 10) E. Graph showing the kupffer cells (Tim4+) ratio in Macrophages (n= 7, 8, 8, 11) and MerTK+ cells in kupffer cells from FACS analysis on day 14. (n=6) F. Graph illustrating the sub-cluster proportions of Mo/MF cells among CD45+ cells. This analysis shows the percentage of activated Mo/MoMFs in Cluster 0 for each group. G. UMAP showing the expression of Ly6C2 and Adgre1 in activated Mo/MFs. H. Paga-velocity analysis image illustrating the differentiation levels of each sub-cluster. The graph shows the directional differentiation of each subcluster over pseudo-time (the right graph displaying the vector values indicating the direction of differentiation.) I. Heatmap of gene expression dynamics for the top 300 genes, selected based on their fit likelihood scores, in each cluster. These genes are ordered along latent time inferred from RNA velocity analysis using scVelo, where fit likelihood represents how well the gene-specific parameters (transcription, splicing, and degradation rates) match the observed data within the dynamic model framework. Each column displays group means with individual data points and error bars with SEM. Statistical significance was determined using one-way ANOVA followed by Tukey’s multiple comparison test (C,D,E). P values indicate significant differences (*p<0.033; **p<0.002; ***p<0.001).

### scHIF1α EVs Accelerate Macrophage Differentiation in Sepsis-induced Liver Injury

During microbial infection, infected cells and Kupffer cells release various signals to recruit other immune cells, particularly monocytes/macrophages, to the liver.^8,30^ Therefore, we hypothesized that scHIF1α EVs might have impacted the accumulated monocytes and macrophages, so we isolated NPCs from the liver and conducted immunological analysis. In the scHIF1α EV group, the proportion of neutrophils, which are known to induce severe inflammatory responses in early sepsis,^3,5^ decreased. Additionally, the proportion of LSECs, which are known to interact with hepatocytes and Kupffer cells induce macrophage differentiation,^13,31^ increased. However, the proportions of Kupffer cells, MoMFs, and monocytes that directly took up the EVs did not show significant differences (Fig 2C). Notably, the proportion of Ly6Clo MoMFs increased (Fig 2D). Ly6C expression is known to decrease as monocytes differentiate into macrophages.^32,33^ Therefore, we hypothesized that these Ly6Clo MoMFs might compensate for the rapidly reduced Kupffer cell population during the early sepsis stage and help restore the Kupffer cell population.^34^ Two weeks after inducing sepsis, we confirmed that the proportion of Kupffer cells and the expression of MerTK^34^ in Kupffer cells were restored in the scHIF1α EV group (Fig 2E). Based on these results, we hypothesized that monocytes/MoMFs that have taken up scHIF1α EVs could promote differentiation into more functional macrophages.

To clarify the mechanisms in macrophage differentiation and functions by scHIF1α EVs, we performed scRNA sequencing on CD45+ cells sorted from liver NPCs 24 hours after LPS treatment. For quality control, we analyzed cells with feature counts between 500 and 8,000, excluded cells with more than 5% mitochondrial genes, and analyzed cells with RNA counts between 1,500 and 60,000. We annotated each cluster based on the expression of representative markers (Extended Fig 3C). Unlike the PBS group, the LPS-treated groups showed a dramatic increase in the proportions of Activated Mo/MoMFs, NeuMos, and Activated Neutrophils. However, these proportions did not significantly differ between the LPS-treated groups (Extended Fig 3D). Despite the lack of significant differences in proportions, previous FACS results indicated changes in macrophage subtypes. Thus, we performed an advanced analysis of Activated Mo/MoMFs to identify subtypes and their functions.

To identify the subtypes of Activated Mo/MoMFs, we divided six clusters based on representative markers, including Cluster 0 (Hmox1, Abca1), Cluster 1 (Ly6c2, Sdf2l1), Cluster 2 (Bcl2, Nr4a1), Cluster 3 (Hist1h1b, Mki67), Cluster 4 (S100a8, S100a9), and Cluster 5 (Hoxa9, Meis1) (Extended Fig 3F). In the scHIF1α EV group, we observed an increase in Cluster 0 and 2 (Fig 2F). Similar to the FACS analysis results, these clusters showed low Ly6c2 expression and high Adgre1 expression, indicating a more differentiated subtype (Fig 2G). Additionally, the expression of Ly6c2 and Itgam was reduced in macrophages of the scHIF1α EV group (Extended Fig 4A). Using scVelo analysis to predict cell type differentiation stages^35^, we identified that these two clusters are the most differentiated macrophages. The overall differentiation trajectory is predicted to progress from cluster 3 to 1, and from cluster 1 to clusters 0 and 2 (Fig 2H, I). Indeed, cluster 2 shows the highest velocity length values (Extended Fig 3D, E). These results indicate that scHIF1α EVs promote the differentiation of recruited Mo/MoMFs.

### scHIF1α EVs Promote Differentiation of Liver Macrophages into Pro-Efferocytic and Pro-Survival MoMFs

To annotate each macrophage subtype, we identified distinctively expressed genes in each cluster and performed Gene Ontology analysis to classify them according to their respective roles. To identify the roles of Cluster 0 and Cluster 2, we examined the genes differentially expressed in each cluster (Fig 3A, B). Cluster 0 exhibited high expression of scavenger receptors^36,37^ and Kupffer cell identity genes^31,34^ (Mertk, Stab1, C1qa, Nr1h3, ID1 and ID3), and was annotated as pro-efferocytic MoMFs. Cluster 2 showed elevated expression of survival genes^34,38^ and JAK-STAT related genes^39,40^ (Bcl2, Gpx1, Bcl6, Bak1, Ifngr1, Il6ra, Jak1and Stat3), and was annotated as pro-survival MoMFs (Fig 3B). Based on GO analysis, we also named the other clusters as cluster 1, identified as Infiltrating CCR2+ MoMFs, cluster 3 as proliferating Mo/MoMFs, cluster 4 as inflamed MoMFs, and cluster 5 as Ribosome-Enriched MFs. (Extended Fig 4B). To effectively distinguish clusters 0 and 2 for in vivo FACS analysis, we selected candidate membrane proteins that were distinctly expressed in each cluster, and had commercially available conjugated antibodies. We chose C3aR1 as the surface protein marker for Cluster 0 and CD9 for Cluster 2. Further experiments were conducted to verify the distinction of these two clusters using FACS analysis (Fig 3C).

**Fig. 3.**
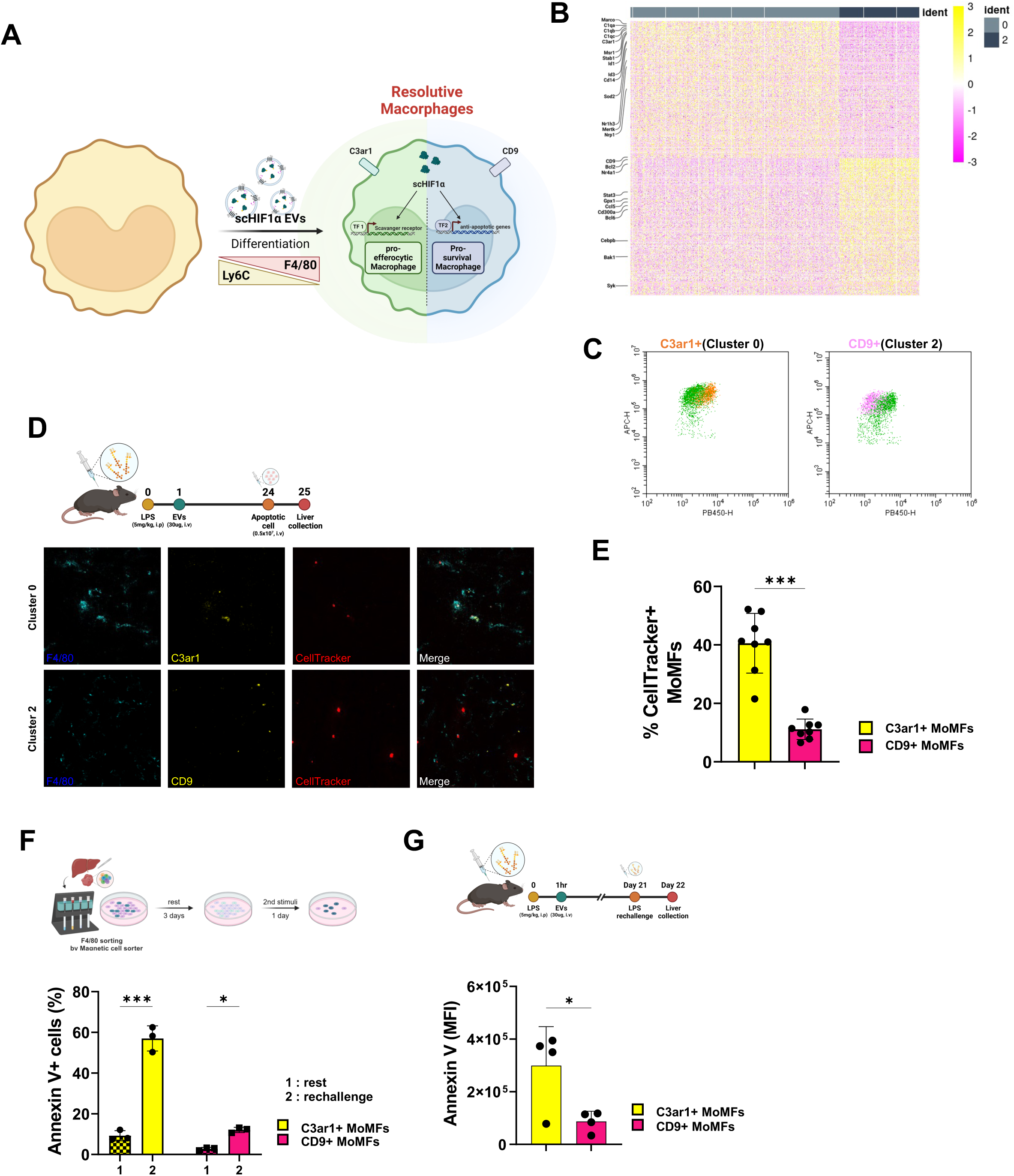
Differentiation into pro-efferocytic and pro-survival macrophages induced by scHIF1α EV. A. Illustration depicting the differentiation of macrophages into pro-efferocytic and pro-survival types induced by scHIF1α EV. B. Heat map based on scRNA-seq data showing differentiated gene expression in clusters 0 and 2. Cluster 0 shows increased expression of scavenger receptors (Marco, C1qa, C1qb, C1qc, C3ar1, Msr1, Stab1, Cd14, Sod2, Mertk, Nrp1) and Kupffer cell imprint transcription factors (Nr1h3, ID1, ID3). Cluster 2 exhibits increased expression of anti-apoptotic genes (Cd9, Bcl2, Nr4a1, Gpx1, Ccl5, Cd300a, Bcl6, Cebpb, Bak1, Syk) and Jak-stat genes (Stat3, Ifngr1, Il6ra, Jak1) C. In vivo FACS analysis showing the expression of C3ar1 (Cluster 0) and CD9 (Cluster 2) in liver MoMFs of LPS-treated mice. D. Immunofluorescence images from in vivo phagocytosis assays of MoMFs in each cluster. Confocal microscopy reveals a higher rate of co-localization between CellTracker and C3ar1+ MoMFs (Cluster 0, pro-efferocytic) compared to CD9+ MoMFs (Cluster 2, pro-survival). E. Graph showing the proportion of CellTracker+ MoMFs as determined by FACS analysis. (n=8) F. Graph depicting the percentage of AnnexinV+ macrophages upon rechallenge with LPS. After sorting liver tissue macrophages in an LPS-induced sepsis model and a 3 day rest, LPS (100 ng/ml) was rechallenged, and the Annexin V+ MoMFs were analyzed. (n=3) G. Graph showing the MFI of AnnexinV+ macrophages upon rechallenge with LPS. After 3 weeks in the LPS-induced sepsis model, LPS (15 mg/kg, i.p) was rechallenged and analyzed. (n=4) Each column displays group means with individual data points and error bars with SEM. Statistical significance was determined using t-test (E,G) and two-way ANOVA followed by Tukey’s multiple comparison test (F). P values indicate significant differences (*p<0.033; **p<0.002; ***p<0.001).

MerTK is known to recognize and phagocytose apoptotic cells by binding to protein S, Gas6, and ptdSer.^41,42^ Marco is a well-known scavenger receptor that phagocytoses unopsonized pathogens.^43^ We hypothesized that Cluster 0, which have high expression of scavenger receptors, would play a role in clearing apoptotic cells during sepsis. To assess the efferocytic capacity of C3aR1+ MoMFs, we intravenously injected CellTracker-labeled apoptotic cells and observed their colocalization with C3aR1+ MoMFs using confocal imaging (Fig 2D). Quantification of the in vivo apoptosis assay via FACS analysis showed that macrophages took up more apoptotic cells than neutrophils, with no significant difference between Kupffer cells and MoMFs (Extended Fig 4D). However, C3aR1+ MoMFs efferocytosed more apoptotic cells than CD9+ MoMFs (Fig 3E). These C3ar1+ MoMFs are similar to Marco+ immunosuppressive macrophages, providing protection in the peri-portal area against commensal-driven liver inflammation.^36^

The expression of Bcl-2 plays a role in protecting macrophages against nitric oxide-induced apoptosis and also known to accumulate at sites of injury and differentiate into tissue remodeling phenotype.^30,44,45^ Additionally, Bcl-2 is known to function as an innate macrophage that can survive for extended periods and perform memory functions.^33,46^ Therefore, we hypothesized that pro-survival MoMFs expressing anti-apoptotic genes would be a death-resistant subtype in inflammatory conditions, potentially inducing endotoxin resistance in secondary infections.^28,44,46,47^ In the sepsis model, we sorted liver NPCs with F4/80 beads, rested cells for three days, and then rechallenged with LPS. The proportion of Annexin V+ cells in C3aR1+ MoMFs significantly increased, whereas the proportion of Annexin V+ cells in CD9+ MoMFs remained relatively low (Fig 3F). Similarly, in in vivo conditions, after rechallenging with a lethal dose of LPS 3 weeks later, it was observed that the Annexin MFI values of CD9+ MoMFs were significantly lower (Fig 3G).

These results suggest that pro-efferocytic macrophages promote the clearance of apoptotic cells, and pro-survival macrophages endure hyperinflammatory conditions to prepare for secondary infections, indicating that these two macrophage types represent a resolutive phenotype to alleviate hyperinflammatory conditions in sepsis (Fig 3A).

### scHIF1α EVs Induce Differentiation of Macrophages Through Upregulation of Nr1h3 and C/ebpβ

To understand the mechanism by which scHIF1α EV induces differentiation into the two types of resolutive macrophages, we examined the differentially expressed RNAs in the scHIF1α EV group using a volcano plot (Fig 4B). Our data analysis identified 8,541 genes that were differentially expressed in the scHIF1α EV group. We focused on genes with a p-value less than 10^-10^. The volcano plot revealed a decrease in Ly6c2 expression, consistent with the FACS data, and an increase in the expression of downstream genes of Nr1h3 (Id3, Jun, Gdf15, Id1, Apobec1, Apoe and Irak2) or C/ebpβ (Atf3, Ccl4, Il1b, Rgs1, Neat1, Dusp2, Clec4e, Nfe2l2, Malat1 and Fnip2) (Fig 4B). Additionally, we confirmed the increased expression of Nr1h3 and C/ebpβ in the scHIF1α EV group (Fig 4C). Nr1h3 is known to be a critical transcription factor for Kupffer cell development and maintenance.^37,48^ Based on the expression of genes related to Kupffer cell imprinting^31,37^ (Nr1h3, Id1 and Id3) and scavenger receptors^34,36,37^ (Mertk, CD163 and Marco) concentrated in pro-efferocytic MoMFs (Extended Fig 4E), we presume that the proportion of recovered Kupffer cells observed after 14 days is derived from these MoMFs (Fig 2E). C/ebpβ is known to be involved in the host’s reorganization of cell lineages in response to bacterial infections^47^ and is a key transcription factor that regulates emergency hematopoiesis by inducing trained immunity in myeloid cells.^46^ Both of these genes are associated with HIF1α. Nr1h3 expression can be upregulated by both HIF1α and the uptake of apoptotic cells.^49,50^ C/ebpβ is directly regulated by Stat3, which is activated by the Jak-Stat signaling pathway in response to interleukins such as IL-6 and IL-1b^39,51^, and Stat3 is also regulated by HIF1α.^40,52^

**Fig. 4.**
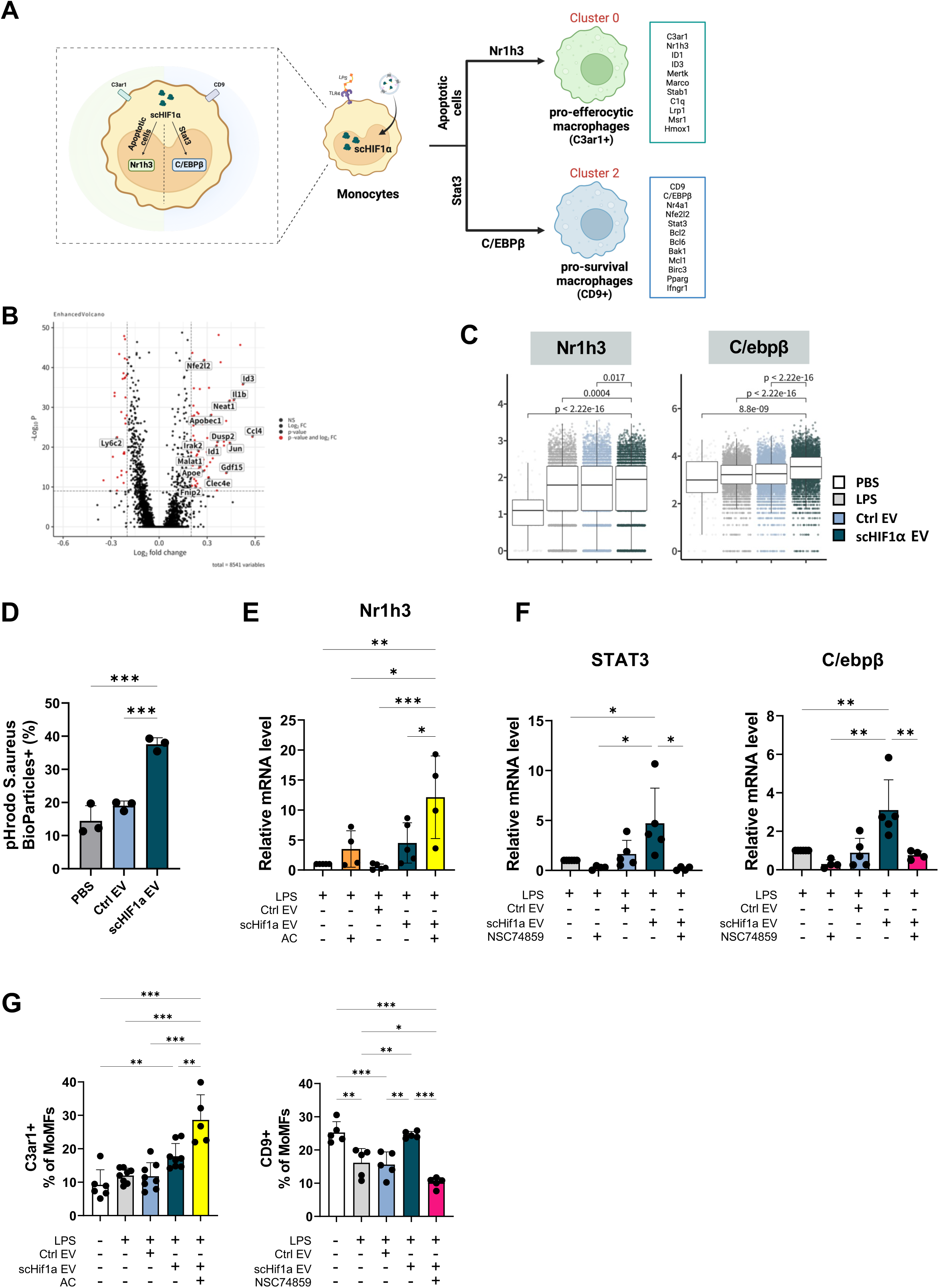
Differentiation of pro-efferocytic MoMFs is Nr1h3-dependent, and pro-survival MoMFs is C/ebpβ-dependent. A. Illustration depicting the molecular mechanisms leading to the differentiation into pro-efferocytic and pro-survival MoMFs. B. Volcano plot showing gene expression changes in MoMFs upon treatment with scHIF1α EVs. Increased expression of Nr1h3 downstream genes (Id3, Jun, Gdf15, Id1, Apobec1, Apoe, Irak2) and C/EBPb downstream genes (Atf3, Ccl4, Il1b, Rgs1, Neat1, Dusp2, Clec4e, Nfe2l2, Malat1, Fnip2). C. Bar plot graph displaying the expression differences of Nr1h3 and C/ebpβ in each group. D. In vitro phagocytosis assay using pHrodo Red S. aureus BioParticles in BMDMs. Differentiated BMDMs were treated with each EVs. After 24 hours, pHrodo Red S. aureus BioParticles were added. E. Graph showing qRT-PCR results of Nr1h3 and C/ebpβ mRNA expression in LPS-treated BMDMs. After treating differentiated BMDMs with LPS, IFNr for 1 hour, each group was treated with EVs. Apoptotic splenocytes were added 1 hour after EV treatment. NSC74859 (Stat3i) was treated 1 hour before LPS treatment. F. Graphs depicting the differentiation of C3ar1+ MoMFs and CD9+ MoMFs through efferocytosis and Stat3-dependent pathways in LPS-induced sepsis model. (left) After 1 hour of LPS (5mg/kg) ip injection, Ctrl or scHIF1α EVs were administered intravenously, followed by the injection of apoptotic splenocytes intravenously, with tissues collected after 1 hour. (right) NSC74859 (Stat3i) 5mg/kg was injected intraperitoneally 0.5 hour after LPS (5mg/kg) injection, followed by Ctrl or scHIF1α EVs intravenously after 0.5 hour, with tissue collection and analysis after 24 hours. G. Graphs depicting the differentiation of C3ar1+ MoMFs and CD9+ MoMFs through efferocytosis and Stat3-dependent pathways in LPS-induced sepsis model. (left) After 1 hour of LPS (5mg/kg) ip injection, Ctrl or scHIF1α EVs were administered intravenously, followed by the injection of apoptotic splenocytes intravenously, with tissues collected after 1 hour. (right) NSC74859 (Stat3i) 5mg/kg was injected intraperitoneally 0.5 hour after LPS (5mg/kg) injection, followed by Ctrl or scHIF1α EVs intravenously after 0.5 hour, with tissue collection and analysis after 24 hours. Each column displays group means with individual data points and error bars with SEM. Statistical significance was determined using one-way ANOVA followed by Tukey’s multiple comparison test (D,E,F,G). P values indicate significant differences (*p<0.033; **p<0.002; ***p<0.001).

Therefore, we aimed to confirm whether scHIF1α EVs regulate the differentiation of clusters by increasing the expression of Nr1h3 and C/ebpβ. First, HIF1α is well known for enhancing the phagocytosis of immune cells,^53,54^ so we hypothesized that scHIF1α EVs could increase phagocytosis in vivo and subsequently upregulate the expression of Nr1h3. To determine whether scHIF1α EV enhances macrophage phagocytosis, we treated BMDMs with pHrodo Red S. aureus BioParticles and observed the uptake after 1 hour. We found that phagocytosis increased in BMDMs treated with scHIF1α EVs (Fig 4D). To confirm whether Nr1h3 is upregulated by scHIF1α EVs, we treated BMDMs with scHIF1α EVs and verified the increase in Nr1h3 expression using RT-PCR. Co-culturing with apoptotic cells led to a synergistic increase in Nr1h3 expression (Fig 4E). Second, it is well known that HIF1α is related to Stat3^40,52^, we hypothesized that scHIF1α EVs might also influence the expression of Stat3 and its downstream target, C/ebpβ. Therefore, treatment with scHIF1α EVs in BMDMs confirmed the increased expression of Stat3 and C/ebpβ using RT-PCR. To determine whether C/ebpβ expression is Stat3-dependent, we treated the cells with the Stat3 inhibitor ^55^ (NSC74859) and observed a decrease in C/ebpβ expression (Fig 4F). In a sepsis model, intravenous injection of apoptotic cells increased the differentiation of C3aR1+ MoMFs, while intraperitoneal injection of the Stat3 inhibitor decreased the differentiation of CD9+ MoMFs (Fig 4G, Extended data 5A). However, the uptake of apoptotic cells did not affect the differentiation of CD9+ MoMFs, nor did the Stat3 inhibitor affect the differentiation of C3aR1+ MoMFs (Extended Fig 5B). Based on these results, we concluded that scHIF1α EVs promoted the efferocytosis, enhancing the differentiation of Nr1h3-dependent pro-efferocytic macrophages, and increased Stat3 expression, promoting the differentiation of C/ebpβ-dependent pro-survival macrophages (Fig 4A).

### scHIF1α EV-Induced Macrophage Zonation in the Periportal Region Enhances Pathogen Clearance and Protects Liver Integrity

Liver tissue, composed of hexagonal units called lobules, exhibits a gradient of blood substances from the portal vein to the central vein, driven by the direction of blood flow.^56^ This structural specificity leads to the distribution of metabolically and immunologically distinct cell populations in each zone.^36,57^ Specifically, during systemic infection and inflammation, pathogens are translocated from the PV to the CV, and the periportal zonation of macrophages facilitates efficient pathogen clearance and minimizes liver damage.^11^

We used spatial transcriptomics to investigate the zonation of pro-efferocytic and pro-survival MoMFs differentiated by scHIF1α EVs during sepsis. First, we confirmed the effective delivery of scHIF1α EVs through the activation of hypoxia-related signaling pathways (Extended Fig 6A). Correlating with scRNA seq data, we observed increased expression of Nr1h3 and C/ebpβ in the scHIF1α EV group (Fig 5A). Additionally, we observed an increase in the activation of the JAK-STAT signaling pathway related to Stat3, the upstream regulator of C/ebpβ (Extended Fig 6A). These results indicate that the findings from scRNA seq are consistent with those observed in spatial transcriptomics.

**Fig 5.**
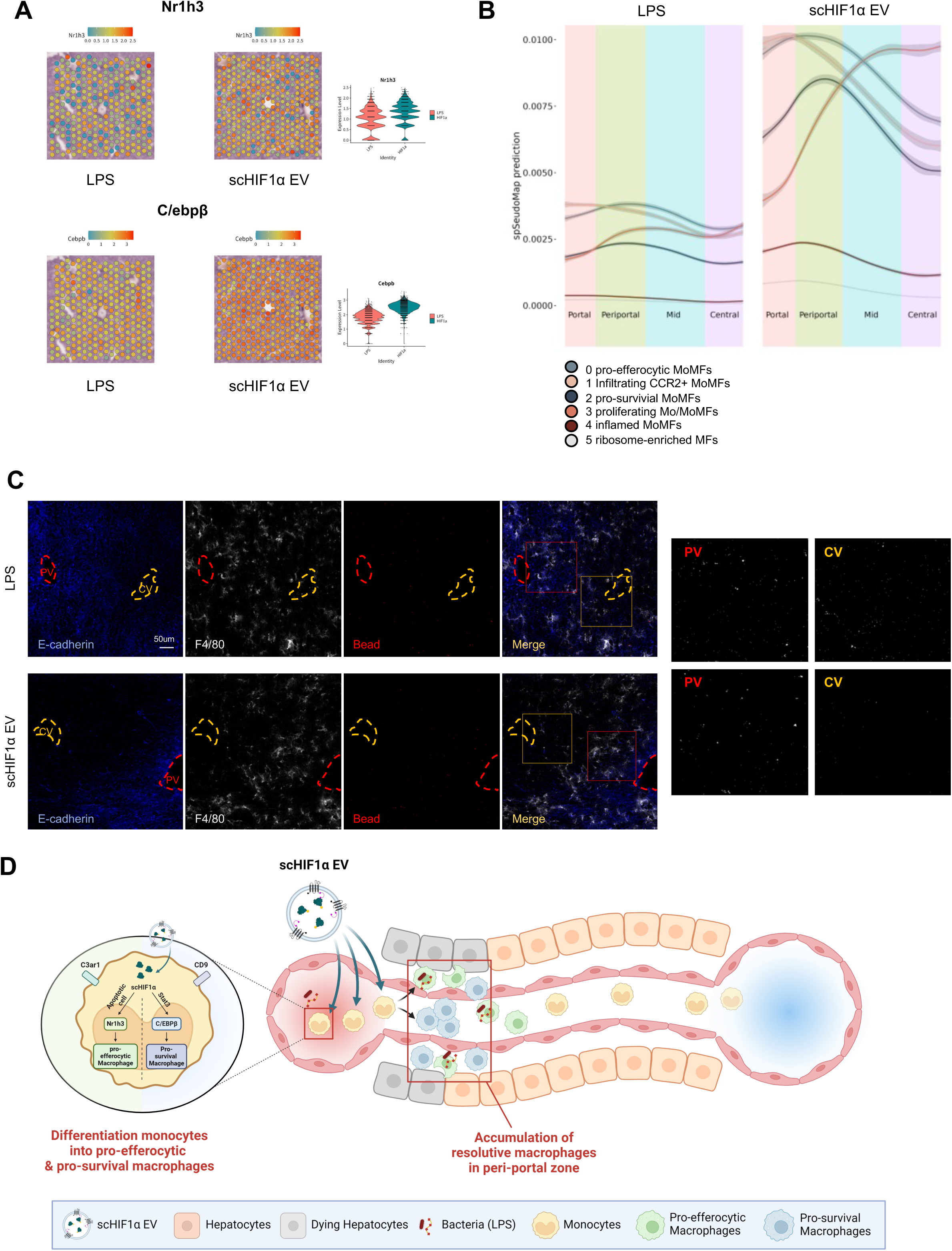
Resolutive MoMFs Form a Wall in the Peri-portal Zone to Prevent the Spread of Damage and Pathogens. A. Liver zonation based on spatial transcriptomic analysis. (left) Cyp2e1, Lect2, Mup11, oat and Scl1a2 represent the central zone (top), Aldh1b1, Apoa4, Cyp2f2, Hsd17b13, Sds represent the portal zone (bottom). (right) Spatial transcriptomics of liver tissue centered on CV, with the upper part showing LPS treatment and the lower part showing treatment with scHIF1α EVs. B. Spatial transcriptomics showing the spatial distribution of each macrophage subtype based on scRNA-seq data, indicating the positioning of macrophages within the liver. C. Immunofluorescence images showing the distribution of macrophages and beads within liver tissue. After 1 hour of LPS (5mg/kg) intraperitoneal injection, scHIF1α EVs were injected intravenously. After 24 hours, pHrodo Red S.aureus BioParticles were injected intravenously, and tissues were collected 0.5 hours later. D. Illustration showing the periportal zonation and physiological functions of Resolutive MoMFs induced by scHIF1α EVs

To distinguish liver zones based on blood flow^56^, we categorized the regions as Portal, Periportal, Mid, and Central based on the expression of PV markers^36,56^ (Aldh1b1, Apoa4, Cyp2f2, Hsd17b13 and Sds) and CV markers^36,56^ (Cyp2e1, Lect2, Mup11, Oat and Slc1a2) (Extended Fig 6B). We obtained spSeudoMap prediction values^58^ to apply the location values of MoMF clusters based on scRNA seq data. We found that LPS treatment disrupted the zonation of all macrophage subtypes. In contrast, in the scHIF1α EV group, pro-efferocytic MoMFs and pro-survival MoMFs maintained their zonation in the Periportal area (Fig 5B). To confirm whether the periportal zonation of pro-efferocytic and pro-survival MoMFs prevents the spread of pathogens, we injected S. aureus BioParticles™ intravenously in a sepsis model and observed bead expansion and macrophage zonation. In the LPS group, macrophage zonation was disrupted, and the beads expanded broadly across the PV-CV region. In the scHIF1α EV group, macrophages accumulated in the periportal zone, and the zonation of these macrophages prevented S. aureus BioParticles™ from spreading to the central vein, leading to their accumulation near the portal vein (Fig 5C). These results suggest that Resolutive MoMFs, specifically pro-efferocytic and pro-survival MoMFs, zonated in the peri-portal region can effectively clear microbes from the bloodstream, preventing infection in the central vein. Consequently, in sepsis, scHIF1α EVs minimize liver damage and promote effective immunological resolution.

## Discussion

Our study emphasizes the pivotal role of HIF1α in reprogramming macrophages during sepsis, presenting a potential therapeutic approach. Delivering the transcription factor HIF1α via EVs significantly reduced severe inflammation and facilitated a transition from an excessively inflammatory environment to a resolutive state. We demonstrated through mechanistic studies that HIF1α EVs induce the differentiation of macrophages into pro-efferocytic and pro-survival MoMFs in an Nr1h3 and C/ebpβ-dependent manner, promoting the zonation of these resolutive macrophages in the periportal region. Additionally, this zonation of resolutive macrophages in the periportal area prevented the spread of infection to the central vein. This strategic localization allowed for effective microbial clearance and protection against secondary infections, suggesting a robust immunological resolution mechanism in sepsis.

Until now, strategies to treat sepsis with excessive immune activation have relied on forcibly suppression of inflammation using high dose glucocorticoids in the early stages. However, simply suppressing inflammation fails to adequately clear infected cells and pathogens, and discontinuing immunosuppression may reduce long-term survival rates due to secondary infections. Therefore, solely suppressing inflammation cannot completely treat sepsis. It is crucial to appropriately maintain necessary inflammatory responses and rapidly deploy resolutive immune cells.

Through scRNA-seq analysis, we discovered that macrophages exhibit a heterogeneous and complex distribution during the initial immune response in sepsis, which cannot be simply categorized as pro-inflammatory or anti-inflammatory. We emphasize that using scHIF1α EVs to induce rapid transitions to resolution among these heterogeneous macrophage populations is essential. Our findings suggest that HIF1α acts as a regulator of functional flexibility in the context of dysregulated inflammatory reactions, allowing macrophages to adapt and respond more effectively to the dynamic demands of the septic environment. This study proposes that leveraging EV-mediated HIF1α delivery offers a novel and promising therapeutic strategy for managing sepsis by precisely modulating immune responses and preventing secondary infections. This research paves the way for future studies to explore and refine approaches for transcription factor delivery, potentially improving survival and recovery outcomes in sepsis patients.

## Method

Detailed information on reagents and Antibodies used in experiments can be found in Supplementary Table 1 and 2.

### Plasmid Construction

All transgenes, except for Flag_CD81_I-CM_Luciferase_myc and Cre recombinase ordered from m.biotech (Gyeonggi-do, Republic of Korea), were obtained by PCR using Phusion High-Fidelity DNA polymerase. The protein constructs utilized in this study were previously described, with codon optimization achieved via the Codon Optimization Tool (Integrated DNA Technologies, USA). The PCR products were cloned into pcDNA3.1(+) Mammalian Expression vector backbone through restriction enzyme cloning and the In-fusion HD Cloning. Plasmids were transformed into TOP10 competent cells.

The scHIF1α sequence, provided by professor Il-Jin Kim at Inha University, was then amplified by PCR. The amplified product was subsequently inserted into the Flag_CD81_I-CM_Luciferase_myc vector using In-fusion cloning to create the Flag_CD81_I-CM_scHIF1α_Myc vector (CD81_I-CM_scHIF1α).We generated point mutations N168A and C169A in the CD81_I-CM_scHIF1α vector to create the CD81_I-CM*_scHIF1α construct and inserted a stop codon after the I-CM sequence to create the CD81_I-CM construct.

The CD81_I-CM_3xNLS_3xsfGFP_2xNLS construct was created by PCR and In-fusion cloning using the pHAGE-TO-nls-st1dCas9-3nls-3XGFP-2nls Vector (#64113, Addgene). The codon-optimized firefly luciferase was then inserted into the 3xsfGFP region via restriction cloning, resulting in the creation of the CD81_I-CM_3xNLS_Luciferase_2xNLS vector.

Lastly, to create CD81_I-CM_Cre, the scHIF1α in CD81_I-CM_scHIF1α was replaced with codon-optimized Cre recombinase using In-fusion cloning.

For the luciferase assay, the VEGF promoter primers were designed based on the reference sequence NC_000006.12. Using the MiniBEST Universal RNA Extraction kit, gDNA was isolated from HEK293T cells. The VEGF promoter was amplified from gDNA using PCR. Subsequently, restriction enzyme sites were added, and the amplified promoter was inserted into the pcDNA3.1 vector, replacing the CMV enhancer and CMV promoter regions.

### Cell culture and EVs Preparation

The HEK293T cells (ATCC, CRL-3216) were cultured in Dulbecco’s modified Eagle’s medium (DMEM) with high glucose, supplemented with 1% Antibiotic-Antimycotic (A.A.) and 10% Fetal Bovine Serum (FBS). Cells were incubated at 37 °C with 5% CO_2_. Cell lines were tested for mycoplasma contamination with Mycostrip^TM^ and maintained free of mycoplasma. At 70-80% confluence, cells were transfected with 20 ug of plasmid DNA and 80 ug of 25 kDa Linear polyethyleneimine (PEI) after the medium change to DMEM with 1% A.A. and 1% Glutamax. After 72 h, the medium was collected and EVs were purified following MISEV guidelines. Cell debris and large extracellular vesicles were removed by serial centrifugation (300 xg for 10 minutes, 2,000 xg for 10 minutes, and 10,000 xg for 30 minutes at 4 °C, Beckman Instruments). The supernatant was circulated through 300 or 500 kDa Spectrum Hollow Fiber Filters using the KrosFlo KR2i TFF System (Repligen). Initially, the supernatant was concentrated 3-fold, dialyzed 8-10 fold with phosphate-buffered saline (PBS) to remove non-EV contaminants, and concentrated to the desired volume. Finally, the supernatant was ultracentrifuged at 150,000 xg for 90 minutes at 4 °C using 45 Ti rotor (Beckman Instruments) after filtration through a 0.45-μm PVDF syringe filter. The sEV pellets were resuspended in PBS.

### EVs Characterization

The presence of scHIF1α and marker proteins was confirmed by Western blot analysis using ChemiDoc (BIO-RAD). The size of EVs was identified using a ZetaView (Particle Metrix), and EVs were visualized by cryogenic transmission electron microscopy (Tecnai F20 G2).

### Comparison of EV Loading Efficiency of scHIF1α Using Different Expression Vectors

The pcDNA3.1_scHIF1α vector and the CD81_I-CM_scHIF1α vector were each transfected into HEK293T cells. To ensure equal transfection efficiency, the pcDNA3.1_mCherry vector was co-transfected at a 1:2 ratio with scHIF1α vectors. After transfection, the medium was collected, and EVs were purified.

Cell lysates were obtained by cell scraping with RIPA buffer. The lysates were vortexed three times at 10-minute intervals at 4°C, and then centrifuged at 13,000 xg for 10 minutes at 4°C to collect the supernatants. The protein concentrations were then measured using a DC Protein Assay kit to ensure equal protein amounts for western blot analysis. Western blot samples were prepared by adding 5x SDS buffer (250mM Tris-Cl (pH 6.8), 5% 2-Mercaptoethanol, 10% SDS, 0.1% Bromophenol Blue, 50% Glycerol) and incubating at 95 °C for 5 minutes.

### Luciferase Assay

After seeding HEK293T cells (3x10^5^ cells) in a 6-well plate, the cell medium was changed to DMEM with 1% A.A. and 1% Glutamax the next day. Cells were transfected with a mixture of 200 ng VEGF promoter luciferase vector and 800 ng PEI. The medium was changed 1 hour post-transfection, and 30 µg of EVs (10 µg/ml) were added. After 30 minutes, the medium was replaced. Luciferase activity was measured 24 hours later using Dual-Luciferase assay kit according to the manufacturer’s Instructions.

### RNA isolation and RT-PCR

EVs were added to HEK293T cells (1 µg/ml) incubated for 24 hours. Total RNAs from the cells were purified using a NucleoSpin RNA kit. cDNA was synthesized using the iScript cDNA Synthesis Kit or QuantiTect Rev.Transcription Kit. RT-PCR (Applied Biosystems StepOne Real-Time PCR System) was performed using PowerUp™ SYBR™ Green Master Mix for qPCR according to the manufacturer’s protocol. Primers used for RT-PCR are listed in Supplementary Table 3. GAPDH was used as the housekeeping gene for HEK293T cells, and r28S was used for BMDMs.

### HeLa cells Confocal Imaging

HeLa cells (Korean Cell Line Bank #10002) were cultured in DMEM high glucose, no phenol red containing 10% FBS, 1% A.A. and 1% Glutamax. After replacing the medium with serum-free medium, CD81_I-CM_3xNLS_3xsfGFP_2xNLS EVs were added at a concentration of 10 µg/ml. Two hours later, the medium was changed. The cells were stained with Hoechst to visualize nuclei and imaged using a Leica TCS SP8 confocal microscope (Leica), employing 40x objective lens and 405 nm and 488 nm lasers. Raw imaging data were analyzed using LAS X office software (Leica).

### Animals

Male C57BL/6N (6–8 weeks old) and Balb/c Nude mice (6–8 weeks old) were purchased from Orient Bio (Gyeonggi-do, Republic of Korea). Ai14 B6.Cg-*Gt(ROSA)26Sor^tm14(CAG-tdTomato)Hze^*/J (JAX stock no. 007914) mice (6–9 weeks old) were provided from Dr. Jinhyun Kim, KIST. All mice were kept in environmentally controlled conditions (23 ± 2 °C, 55 ± 10% relative humidity, and a 12-hour light/dark cycle). The mice were provided with unrestricted access to food and water at a specific-pathogen-free animal facility at the Korea Institute of Science and Technology (KIST; Seoul, Republic of Korea). All animal experiment protocols were performed in accordance with the guidelines of the Institutional Animal Care and Use Committee, KIST (approval no. KIST-IACUC-2023-002-4).

### Mouse Liver NPCs Isolation

Livers from C57BL/6 mice were collected and processed using a gentleMACS Dissociator (Miltenyi Biotec) in DMEM with Collagenase Type II (200 µg/ml) under the ’37c_m_LIDK_1’ program to ate a single-cell suspension. After passing through a 40 μm cell strainer, the flow-through was centrifuged at 50 ×g for 3 minutes at 4°C to collect the supernatant. The supernatant was then centrifuged at 150 ×g for 3 minutes to pellet the non-parenchymal cells (NPCs). The NPC pellet was subjected to RBC lysis using RBC Lysis Buffer at 4°C for 5 minutes. After lysis, the cells were washed with DPBS and centrifuged.

### Ex vivo Angiogenesis Cytokine Assay

Macrophages were sorted from isolated NPCs using Anti-F4/80 Micro-Beads. The sorted macrophages (3×10^5^ cells/well) were seeded in a 96-well round-bottom plate. After 24 hours of incubation, the cells were treated with EVs (1 µg/well). The supernatant was collected 24 hours post-treatment and analyzed using a Proteome Profiler Mouse Angiogenesis Array Kit.

### In Vivo Matrigel Plug Angiogenesis Assay

Matrigel® Matrix High Concentration was mixed with EVs at a 3:1 ratio, ensuring each Balb/c nude mouse received 100 µg of EVs. The mixture was injected subcutaneously into each mouse. After one week, the mice were sacrificed. The Matrigel plugs were then collected and photographed for analysis.

### Survival Study

C57BL/6 mice were administered LPS at 15 mg/kg via intraperitoneal injection to induce sepsis. One hour later, EVs were administered intravenously at a dose of 50 ug (0.5 µg/ml). The mice were monitored for survival over a period of 3 days.

### LPS induced Liver Damage Model

C57BL/6 mice were injected intraperitoneally with 5 mg/kg LPS to induce liver injury. One hour later, EVs were administered intravenously at a dose of 30 µg (0.3 µg/µl). After 24 hours, the mice were sacrificed for subsequent analysis.

### Assessing Liver Damage in Sepsis-Induced Liver Injury

The following experiments were conducted after inducing LPS-induced damage in mice. Serum was collected from the orbital sinus using Micro-Hematocrit Capillary Tubes, incubated at room temperature for 6 hours, and centrifuged at 2000 ×g for 20 minutes at 4°C. Serum samples were prepared for biochemical analyses, including the measurement of AST/ALT (DK Korea, Seoul, Republic of Korea), Mouse TNF-alpha, Mouse IL-6, and Proteome Profiler Mouse Cytokine Array using specific assay kits.

Liver NPCs were isolated and incubated with antibodies at 4°C for 1 hour. The cells were then washed with DPBS and centrifuged at 3000 rpm for 3 minutes at 4°C. Analysis was performed using CytoFlex Flow Cytometer (Beckman).

Macrophages were separated from NPCs using Anti-F4/80 Micro-Beads and seeded at 10^5^ cells in RPMI with 10% FBS and 1% A.A.. After 48 hours, the supernatant was collected for Proteome Profiler Mouse Cytokine Array Kit.

A TUNEL assay (DK Korea, Seoul, Republic of Korea) was conducted on formalin-fixed liver tissue to assess cell apoptosis.

### In Vivo Biodistribution Profile of EVs

EVs were labeled with 10 µM Cyanine5 NHS ester and purified using a Zeba Spin Desalting Columns. C57BL/6 mice were injected intraperitoneally with 5 mg/kg LPS. One hour later, 100 µg of Cy5-EVs with same fluorescent intensity were injected intravenously. The mice were sacrificed 1hour post-injection. The tissues were then harvested and analyzed for biodistribution using the IVIS Spectrum System (Caliper Life Sciences). For FACS analysis, livers were collected 5 minutes post-injection.

Ai14 mice were injected with 5 mg/kg LPS. One hour later, a mixture of 30 µg of scHIF1α EVs and 30 µg of Cre loaded EVs, was injected intravenously. Mice were sacrificed on 5 Days after, and tissues were fixed with 4% formalin. After this step, the tissues were sent to PrismCDX (Gyeonggi-do, Republic of Korea) for Multiplex immunohistochemistry analysis.

### Confocal Fluorescence Imaging of Liver Cryosection

Frozen Tissues were cryo-sectioned at 10 µm thickness using Cryocut Microtome (Leica CM1860) and mounted on SuperFrost Plus Adhesion slides. Sections were washed with PBS, blocked with 3% Bovine Serum Albumin (BSA) in PBS for 1 hour, and incubated with antibodies in 1% BSA in PBS for 1 hour at RT. After washing with PBS, a Fluoromount-G was applied, and cover glass was placed for Confocal imaging. Imaging was performed using a 10x objective lens and 405 nm, 488 nm, 514 nm, and 644 nm lasers.

### Single Cell RNA sequencing Sample Preparation and Analysis

NPCs were isolated from a pooled group of LPS-induced damage model mice (n=10) 24 hours later and sorted using CD45 MicroBeads. The single-cell RNA sequencing (scRNA-seq) was outsourced to Macrogen (Republic of Korea). Libraries were prepared using the Chromium Next GEM Single Cell 3’ RNA library v3.1 platform (10x Genomics).

Cell suspensions were partitioned into nanoliter-scale Gel Beads-in-emulsion (GEMs) with Single Cell 3’ v3 Gel Beads containing barcoded oligonucleotides. RNA transcripts from single cells were uniquely barcoded and reverse-transcribed within droplets, producing full-length cDNA, which was enriched through PCR. Libraries were sequenced on the Illumina system, with paired-end reads. Data were processed using the Cell Ranger 7.0.1 pipeline, including demultiplexing (mkfastq), alignment, filtering, barcode counting, and UMI counting (count). Quality control was performed with FastQC v0.11.7.

Single-cell RNA sequencing data were processed using R (v.4.4.0) and Seurat (v.4.3.0). Cells were filtered based on the following criteria: unique feature counts between 500 and 8,000, mitochondrial gene expression below 5%, and RNA counts between 1500 and 60,000. (Ref. 3) The filtered cell data were normalized using the SCTransform, regressing out percent.mito and nCount_RNA, with 3,000 variable features and memory conservation enabled. Dimensional reduction was performed using 50 principal components (PCs).

Uniform manifold approximation and projection (UMAP) was executed for the first 10 PCs. The FindNeighbor was employed for unsupervised clustering, with a resolution parameter set at 0.12. Activated Mo/MoMFs cluster was further subset and reanalyzed using the FindNeighbor with a resolution of 0.3.

Differentially expressed genes were identified using the FindAllMarkers (min.pct = 0.25, logfc.threshold = 0.25), resulting in a list of characteristic genes for each cluster. The significant shared GO terms for biological processes were identified using the PANTHER Classification System (www.pantherdb.org). The data was then visualized using GraphPad Prism (V10.1.2). Additionally, the gene lists were filtered (p_val_adj < 10^-8^, pct.1 > 0.3) to select genes for creating heatmaps with the ComplexHeatmap or dittoSeq.

Volcano plots were generated using gene lists showing significant differences in HIF1α EVs compared to LPS and Ctrl EVs. These differences were identified using the FindMarkers (logfc.threshold = 0, min.pct = 0.1). The volcano plots were then visualized with EnhancedVolcano. Other scRNA-seq analysis datas were further visualized using dittoSeq and ggplot2.

We conducted Trajectory Analysis by referencing the scVelo tutorial (https://scvelo.readthedocs.io/en/stable/). The analysis focused on the following clusters: Cluster 0 (Pro-efferocytic MoMFs), Cluster 1 (Infiltrating CCR2+ MoMFs), Cluster 2 (Pro-Survival MoMFs), and Cluster 3 (Proliferating Mo/MoMFs). The scVelo analysis was performed using Visual Studio Code with Python (v3.8.16).

### Visium

The liver was harvested 24 hours post-LPS induction in the LPS-induced damage model mice, and Formalin-Fixed Paraffin-Embedded (FFPE) samples were prepared. These samples were sent to Portrai (Seoul, Republic of Korea) for visium analysis. The data processing was performed using the SpaceRanger (v.2.1.0) pipeline, and all subsequent steps were carried out with the Seurat (v4.3.0) pipeline in R (v4.3.1). Spots with fewer than 5,000 or more than 50,000 detected transcripts, or those with fewer than 2,000 genes, were excluded from the analysis. The SCTransform function was used for gene identification and scaling, followed by analysis using spSeudoMap and STopover algorithms.

### Apoptotic Splenocytes Preparation

Spleens were collected from C57BL/6 mice. The cells were processed using the gentleMACS Dissociator ‘Spleen_01’ protocol and passed through a 0.4 µm strainer. Following centrifugation at 450 rpm for 10 minutes, the supernatant was removed. The cells were seeded in a Petri dish after RBC lysis. The cells were then exposed to UV light for 90 minutes, stained with CellTracker Deep Red Dye at a concentration of 5 µM, and incubated at 37°C for 20 minutes. After centrifugation at 3000 rpm for 3 minutes, cell counting was performed using FACS.

### In Virto Phagocytosis Assay

BM was obtained from C57BL/6 mice and differentiated into BMDMs. BMDM differentiation and maintenance followed previously established protocols. **(Ref. 4)** BMDMs were seeded at 2*10^5 cells per Falcon dish and incubated for one day before treating with EVs (3 µg/ml). After another day, 10 µg of beads were added, and cells were collected for FACS analysis one hour later.

### In Vivo Phagocytosis Assay

In the LPS-induced damage model, CellTracker Deep Red Dye labeled 0.5 × 10^7^ apoptotic splenocytes were injected intravenously 90 minutes before the mice were sacrificed. Tissues were preserved in OCT compound and frozen at -80°C for subsequent confocal imaging.

### Ex Vivo Macrophage Survival to LPS Rechallenge

Macrophages were sorted from these NPCs using F4/80 sorting beads and 5 × 10^4^ cells seeded onto falcon dish well in RPMI. After 3 days, the cells were rechallenged with LPS (100 ng/ml). One day after the LPS rechallenge, the cells were centrifuged at 150 ×g for 3 minutes, and the pellet was resuspended in 1x Annexin binding buffer (Invitrogen). The samples were incubated with Annexin V for 1 hour at 4°C. Following incubation, the samples were analyzed using FACS.

### In Vivo Macrophage Survival to LPS Rechallenge

In the LPS-induced liver damage model, C57BL/6 or Ai14 mice were administered EVs and allowed a rest period of 3 weeks. Following this, mice were rechallenged with LPS (15 mg/kg) intraperitoneally. Four days after the LPS injection, liver NPCs were isolated and analyzed using FACS.

### In vitro and in vivo assay for Efferocytosis and STAT3-Dependent Macrophage Differentiation

To mimic the LPS-induced damage model, BMDMs were treated with 100 ng/ml LPS and 100 ng/ml INF-γ. One hour later, apoptotic Splenocytes (5 × 10^5^ cells) and NSC74859 (100 µM) were added. After another hour, the medium was replaced, and EVs (3 µg/ml) were added. RNA was then extracted and analyzed via RT-PCR.

In the LPS-induced damage model, 0.5 × 10^7^ apoptotic splenocytes were injected intravenously one hour before the mice were sacrificed, and tissues were collected. In a separate set of experiments, 30 minutes after LPS administration, NSC74859 (5 mg/kg) was administered intraperitoneally. Following another 30 minutes, EVs were injected intravenously. Tissues were collected and analyzed after 24 hours.

### pHrodo Red S. aureus BioParticles Mediated In Vivo Phagocytosis Assay for Macrophage Zonation

In the LPS-induced damage model, 40 µg of pHrodo Red S. aureus BioParticles were administered intravenously 24 hours post-LPS injection. The mice were sacrificed 30 minutes post-administration for further analysis. The tissues were preserved by embedding in OCT compound and stored at -80°C for later confocal imaging.

### Statistics

Statistical analysis was conducted using GraphPad Prism 10. Comparisons between two groups were performed using a two-tailed unpaired Student’s t-test. Comparisons among multiple groups were evaluated using one-way analysis of variance (ANOVA) followed by Tukey’s post-hoc test. For multiple parameter samples, a two-way ANOVA with Sidak’s post-hoc test was used. The survival rate was analyzed with Kaplan–Meier log-rank analysis, and proteomic analysis data were assessed using Pearson’s correlation coefficient. Data are presented as mean ± SD. Statistical significance was denoted as *p < 0.033, **p < 0.002, and ***p < 0.001.

## Acknowledge

This research was supported by the funding sources below. National Research Council of Science and Technology (NST) grant 2017R1A3B1023418 (ISK) and 2021R1A2C1095046 (CJ). Korea Health Technology R&D Project HI22C1501 (ISK) and RS-2023-00266133 (CJ). KU-KIST School Project (ISK). KIST Institutional Program 2V09840-23-P038 (ISK) and 2E33132 (CJ). The grant Nos. 2021-0-02076 and 2023-00262155 (IITP) funded by the Korea government (the Ministry of Science and ICT) as well as the Global Mobility project in KIST. Cryo-TEM imaging (FEI Tecnai F20 G2) was performed at the KIST Advanced Analysis Center (Seoul, Republic of Korea). The visium analysis using spSeudoMap and STopover algorithms was conducted by Portrai (Seoul, Republic of Korea).

## Author Contributions

C.J. and I.-S. K. conceived the study. Y.L. and J.G. designed the experiments. I.K. and J.P. provided the HIF1α plasmid. Y.L. and J.G. performed the plasmid cloning. Y.L., J.G., and S.C. conducted the cell culture and EV purification. Y.L. and J.G. carried out the in vitro assays. Y.L. and J.G. performed the confocal imaging. Y.L., J.G., and S.C. conducted the in vivo assays. Y.L., J.G., and S.K. wrote the code and analyzed the scRNAseq and visium data. Y.L. and J.G. wrote the manuscript. G.N., C.J., and I.-S. K. contributed to the project design.

---

**Extended Figure 1.**
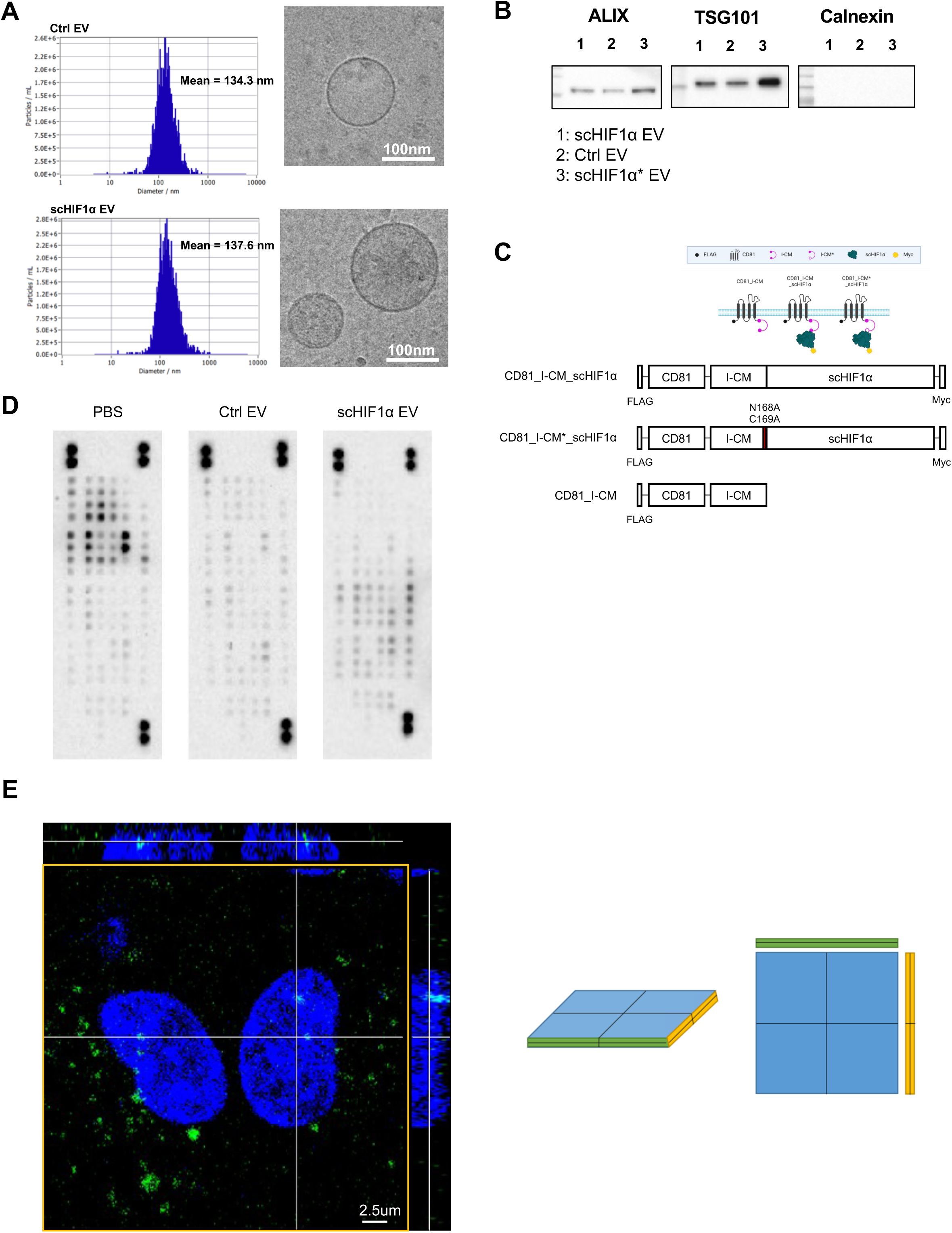
EV Characterization and Platform Construct. A. Characterization of EV size and particle numbers using ZetaView and cryo-TEM. B. Characterization of EV markers using Western blot. C. Illustrations and structures for verifying the cleavage effect of pH-dependent mini intein: CD81_I-CM_scHIF1α (top), CD81*_*I-CM*_scHIF1α (middle), CD81_I-CM (bottom). D. Angiogenesis cytokine array in ex vivo F4/80 sorted macrophages from liver. E. Confirmation of nucleus translocation of CD81_I-CM_3xNLS_3xsfGFP_2xNLS EV via confocal imaging.

**Extended Figure 2.**
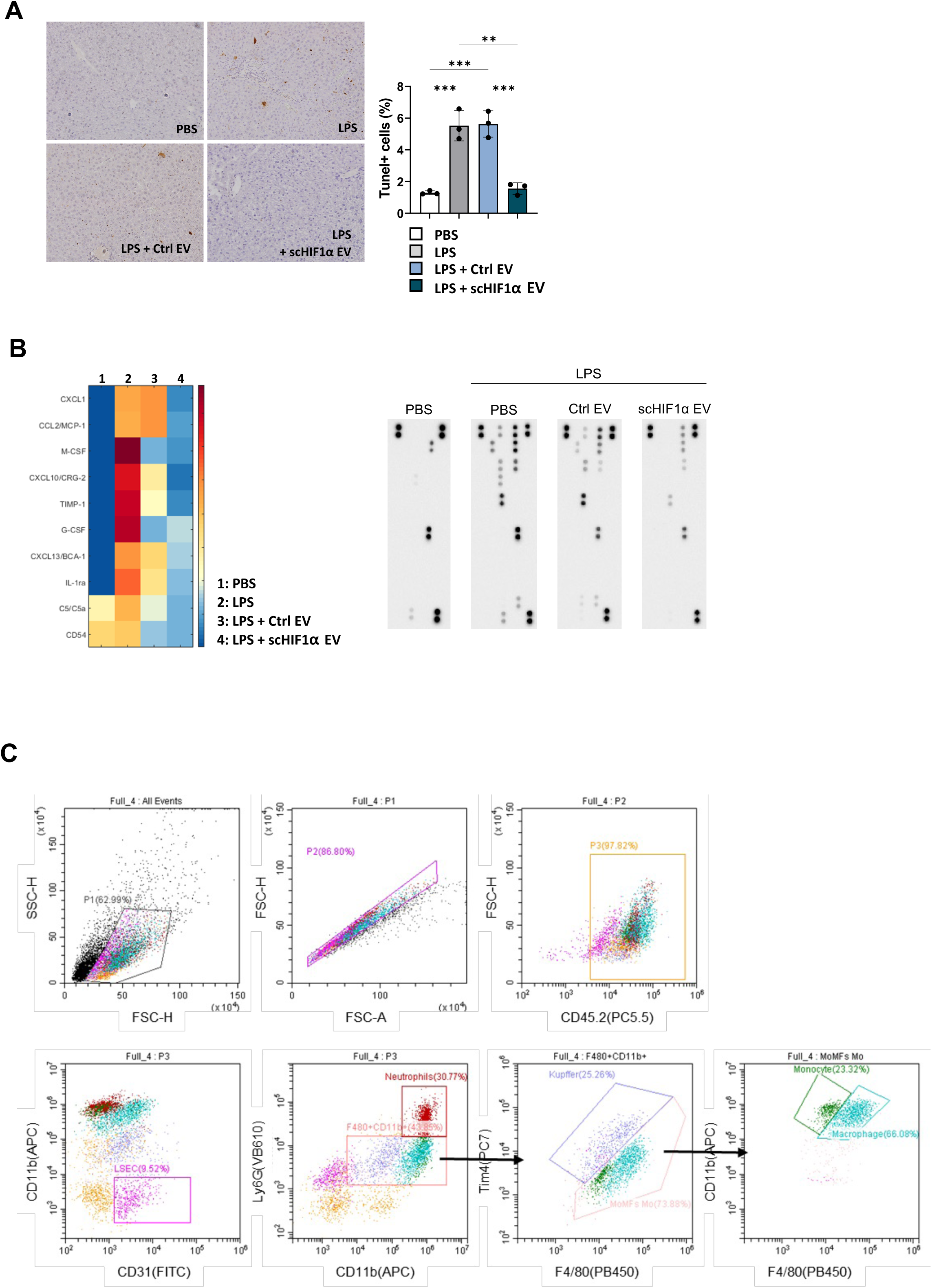
In vivo Efficacy of scHIF1α EV. A. Detection of apoptotic cells in sepsis-induced liver injury mice using the TUNEL assay. The left image shows liver tissue stained by the TUNEL method, the right graph represents the ratio of TUNEL-positive cells to total cells (n=3). B. Serum cytokine array from mice with sepsis-induced liver injury. A heatmap was generated using serum samples from each mouse (n=3), with total intensity normalized to 1 for all four groups to display fold changes. C. In vivo FACS gating strategy. Each column displays group means with individual data points and error bars with SEM. Statistical significance was determined using one-way ANOVA followed by Tukey’s multiple comparison test (A). P values indicate significant differences (*p<0.033; **p<0.002; ***p<0.001).

**Extended Figure 3.**
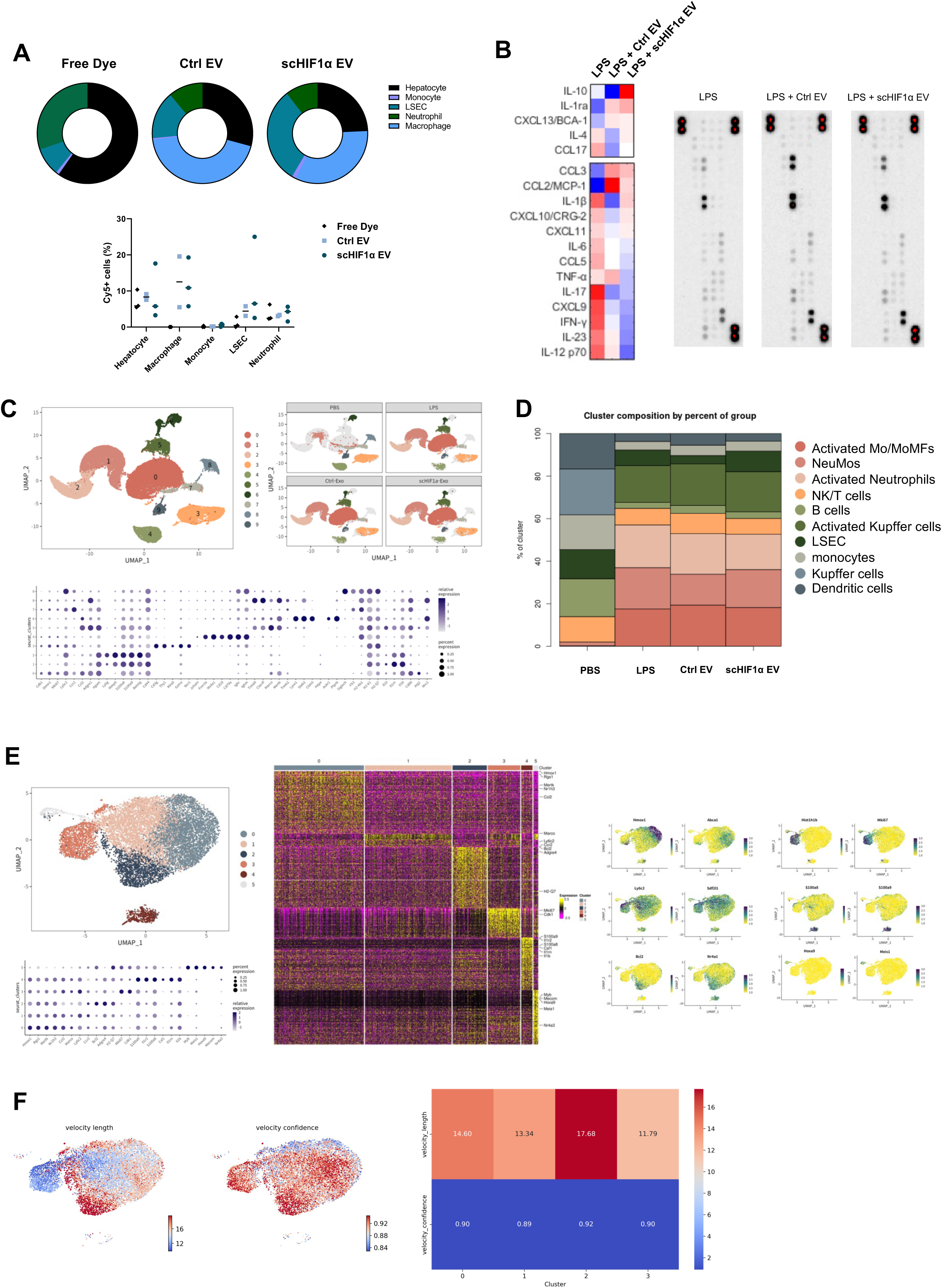
Identification of Immune Cells Uptaking scHIF1α EVs in the Liver and Basic Analysis of Total CD45+ Clusters and Activated Mo/MoMFs Clusters via scRNA seq. A. Analysis of immune cells uptaking Cy5+ labeled EVs. (n = 3, 2, 3) B. Immune suppressive cytokine secretion by macrophages sorted from the liver using F4/80+ beads and treated with scHIF1α EVs. C. Cluster annotation of total immune cells via scRNA seq analysis. D. Proportion analysis of total immune cells via scRNA seq analysis. E. Identification of subcluster markers for activated Mo/MoMFs via scRNA seq analysis. F. Heatmaps of velocity length which indicates the speed or rate of differentiation, and velocity confidence which indicates the coherence of the vector field. The table shows the average values of velocity length and velocity confidence for each cluster. (left) Contour data showing the higher cell density expressing genes related to differentiation in pro-efferocytic MoMFs and pro-survival MoMFs compared to other clusters. These data were analyzed using scRNAseq analysis.(right)

**Extended Figure 4.**
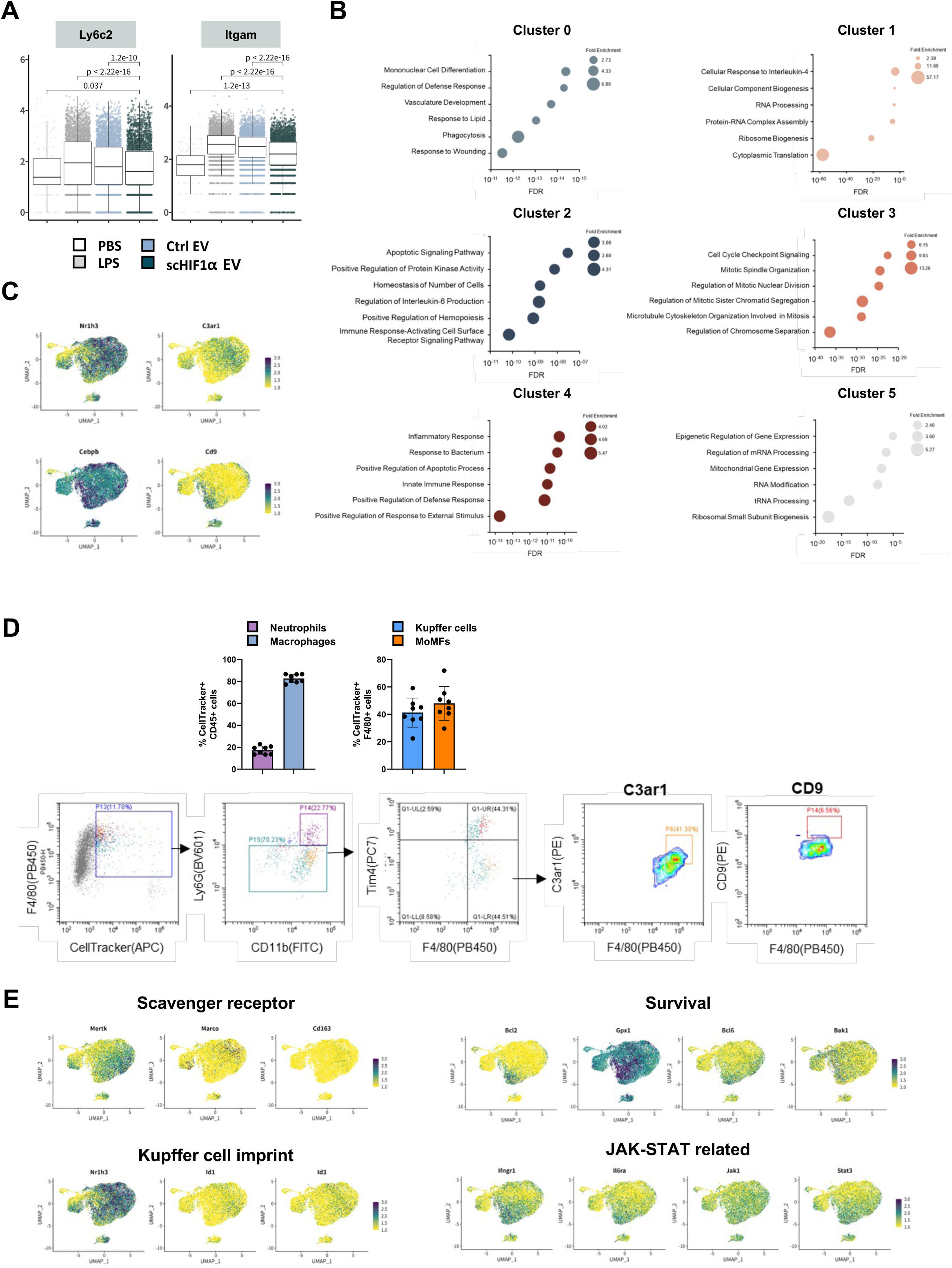
Subcluster Annotation of Activated Mo/MoMFs Clusters and Gene Expression Analysis of Clusters 0 and 2. A. Analysis of Ly6c2 and Itgam expression in activated Mo/MoMFs across different groups. The data were analyzed using R with the ggpubr package, and statistical significance was determined using the Wilcoxon rank-sum test. B. Gene ontology analysis based on distinct RNA expression in each subcluster. C. Increased expression of transcription factors and membrane protein markers in clusters 0 and 2. D. FACS gating strategy for in vivo phagocytosis analysis. (n = 8) E. RNA expression patterns of scavenger receptors(Mertk, Marco, CD163), Kupffer cell imprint (Nr1h3, Id1, Id3), survival markers(Bcl2, Gpx1, Bcl6, Bak1), and Jak-STAT(Ifngr1, Il6ra, Jak1, Stat3) related genes showing distinct expression in clusters 0 and 2.

**Extended Figure 5.**
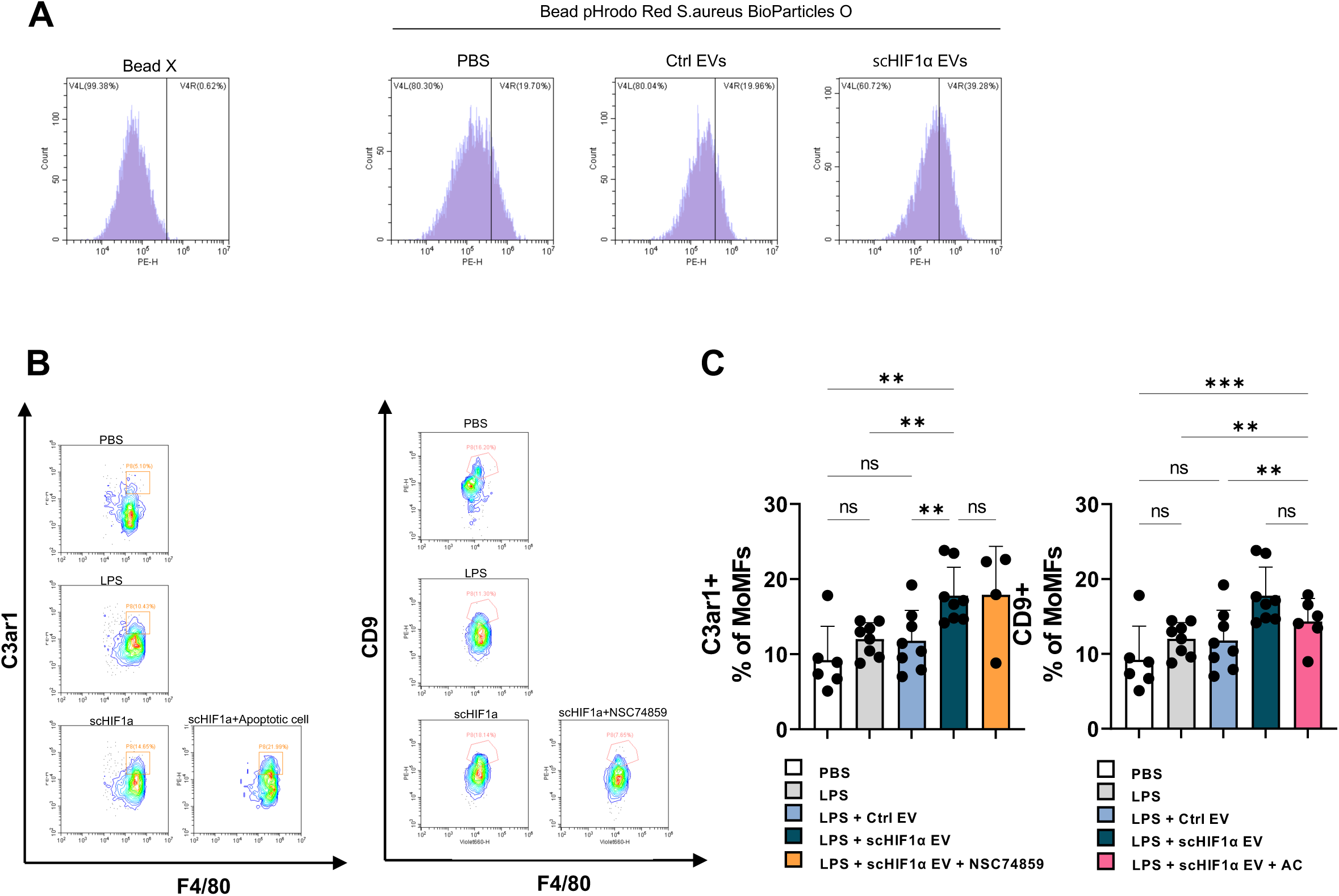
Analysis of Nr1h3 and C/ebpβ Dependent MoMFs Differentiation. A. Relative MFI values in an in vitro phagocytosis assay after treating BMDMs with scHIF1α EVs. B. Comparison of the frequency of C3ar1 and CD9 positive MoMFs, markers of clusters 0 and 2. C. Analysis of the differentiation ratio of STAT3-dependent C3ar1+ MoMFs (n = 6, 8, 8, 8, 4) and apoptotic cell-dependent CD9+ MoMFs (n = 6, 8, 8, 8, 6). Each column displays group means with individual data points and error bars with SEM. Statistical significance was determined using one-way ANOVA followed by Tukey’s multiple comparison test ©. P values indicate significant differences (*p<0.033; **p<0.002; ***p<0.001).

**Extended Figure 6.**
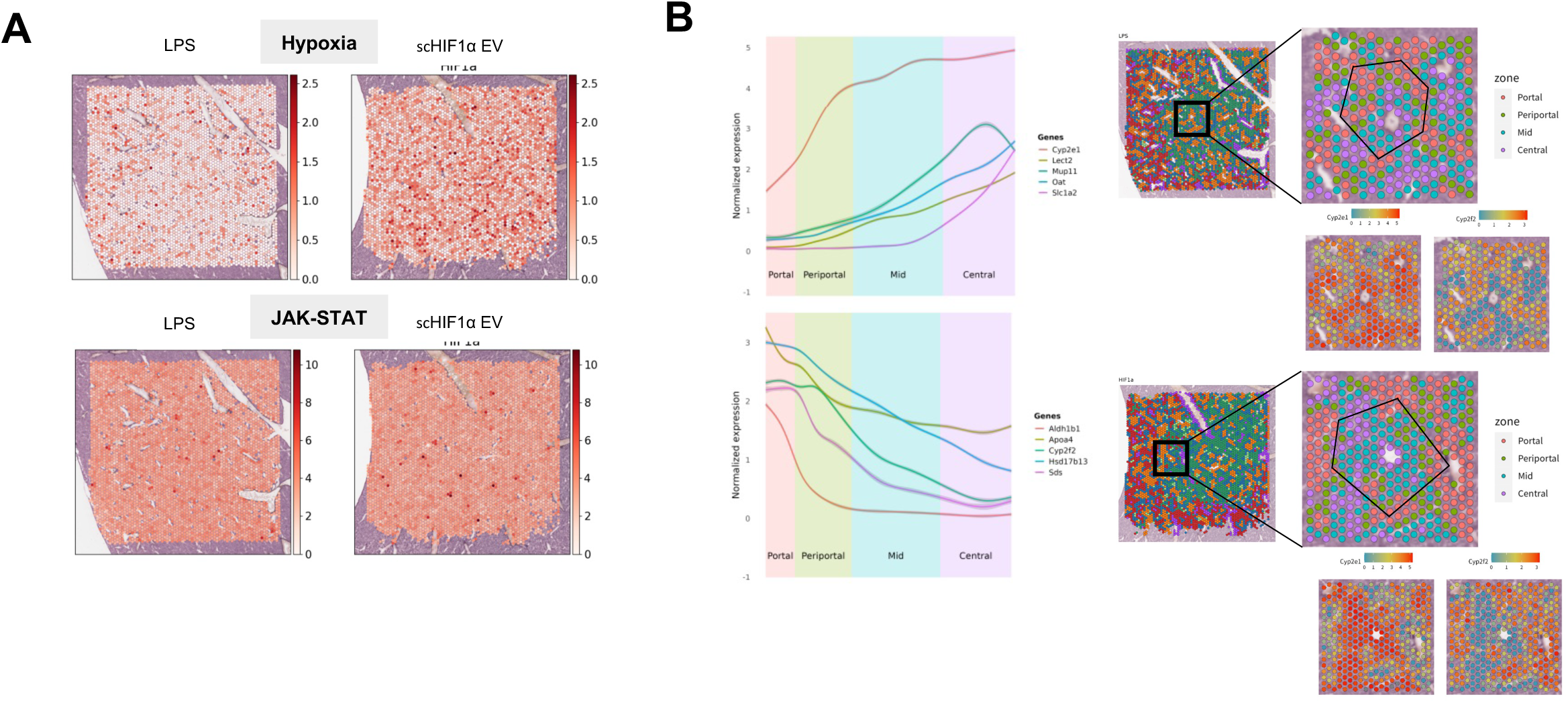
Basic Analysis of Visium and Comparison of RNA Expression Across Zones. A. Analysis showing RNA expression related to hypoxia and JAK-STAT signaling. B. Expression ratio of markers differentially expressed in portal and central zones, distinguishing the respective zones.

