## Supplemental Data for "Extracellular Vesicle-Mediated Delivery of HIF1α Reprograms Macrophages for resolutive response in Sepsis"

**Affiliations**

Supplement Table 1. Reagents used in these experiments

| Product | Vender | Catalog # | Usage |
| --- | --- | --- | --- |
| Phusion™ High-Fidelity DNA Polymerase | Thermo Scientific | F503L | Cloning |
| In-Fusion® HD Cloning Kit | Clontech | 639650 | Cloning |
| MiniBEST Universal RNA Extraction kit | Takara | 9767A | gDNA extraction |
| Cell Culture Dish 150 x 20 mm | Sarstedt | 83.3903 | Cell Culture |
| Cell Culture Dish 90 x 20 mm | SPL | 20100 | Cell Culture |
| Petri Dish | SPL | 10090 | Cell Culture |
| Confocal dish | SPL | 101350 | Cell Culture |
| Falcon® 35 mm Not TC-treated Easy-Grip Style Bacteriological Petri Dish | Corning | 351008 | Cell Culture |
| 6-well Clear TC-treated Multiple Well Plates | Corning | 3516 | Cell Culture |
| 96 well cell culture plate | Corning | 34296 | Cell Culture |
| Dulbecco's Modified Eagle Medium (DMEM) with high glucose | HyClone | SH30243.01 | Cell Culture |
| Dulbecco's Phosphate Buffered Saline (DPBS) | WELGENE | LB001-02 | Cell Culture |
| Antibiotic-Antimycotic (100X) | Gibco | 15240062 | Cell Culture |
| Fetal Bovine Serum, qualified, Canada | Gibco | 12483020 | Cell Culture |
| Mycostrip^TM^ | InvivoGen | rep-mysnc-100 | Mycoplasma Detection |
| Polyethylenimine, Linear, MW 25000, Transfection Grade (PEI 25K™) | polysciences | 23966-1 | Transfection |
| GlutaMAX™ Supplement | Gibco | 35050061 | Cell Culture |
| MIDIKROS 41.5CM 500K MPES 0.5MM FLL X FLL 1/PK | Repligen | D04-E500-05-N | sEV Purification |
| MINIKROS SAMPLER 41.5CM 300K MPES 0.5MM 3/4TC X 3/4TC | Repligen | S04-E300-05-N | sEV Purification |
| MINIKROS SAMPLER 65CM 300K MPES 0.5MM 3/4TC X 3/4TC | Repligen | S06-E300-05-N | sEV Purification |
| 10X PBS | BIOSESANG | HP2007-1 | sEV Purification |
| Millex-HV Syringe Filter Unit, 0.45 µm, PVDF | Millipore | SLHVR33RB | sEV Purification |
| Protein Assay Reagent A | BIO-RAD | 5000113 | Protein Concentration Quantification |
| Protein Assay Reagent B | BIO-RAD | 5000114 | Protein Concentration Quantification |
| Protein Assay Reagent S | BIO-RAD | 5000115 | Protein Concentration Quantification |
| 4-20% Mini-PROTEAN TGX Precast Protein Gels | BIO-RAD | 4561096 | Western Blot |
| Trans-Blot Turbo RTA Mini 0.2 µm Nitrocellulose Transfer Kit | BIO-RAD | 1704270 | Western Blot |
| Clarity Western ECL Substrate | BIO-RAD | 1705061 | Western Blot |
| 100micron A4 OHP Film | jongienara | AME00001 | Western Blot |
| RIPA Buffer (10X) | Cell Signaling TECHNOLOGY | 9806 | Cell Lysate Preparation |
| Polystyrene Beads | Thermo Scientific | 3090A | NTA |
| NucleoSpin RNA kit | Macherey-Nagel | MN740955.250 | RT-PCR |
| iScript™ cDNA Synthesis Kit | BIO-RAD | 1708891 | RT-PCR |
| QuantiTect Rev. Transcription Kit | QIAGEN | 205311 | RT-PCR |
| PowerUp™ SYBR™ Green Master Mix for qPCR | Applied Biosystems | A25741 | RT-PCR |
| 96-well White Flat Bottom Polystyrene Not Treated Microplate | Corning | 3912 | Luciferase Assay |
| Dual-Luciferase® Reporter Assay System | Promega | E1680 | Luciferase Assay |
| DMEM, high glucose, no glutamine, no phenol red | Gibco | 31053028 | Confocal Imaging |
| Hoechst 33342 | Invitrogen | H3570 | Confocal Imaging |
| Collagenase Type II | Gibco | 17101015 | NPCs Isolation |
| gentleMACS C Tube | Miltenyi Biotec | 130-096334 | NPCs Isolation |
| Cell Strainer 40 μm | SPL | 93040 | NPCs Isolation |
| RBC Lysis Buffer (10X) | BioLegend | 420301 | NPCs Isolation |
| Anti-F4/80 MicroBeads UltraPure, mouse | Miltenyi Biotec | 130-110-443 | Macrophage Sorting |
| Proteome Profiler Mouse Angiogenesis Array Kit | R&D Systems | ARY015 | Angiogenesis Cytokine Assay |
| Matrigel® Matrix High Concentration | Corning | CLS354263 | In Vivo Angiogenesis Assay |
| Lipopolysaccharides from *Escherichia coli* O111:B4 | Sigma-Aldrich | L2630-25mg | In Vivo Animal Experiments |
| Micro-Hematocrit Capillary Tubes | CHASE | HCH-41A2502 | Blood Collection |
| Mouse TNF-alpha ELISA Kit | R&D Systems | MTA00B | Cytokine Quantification |
| Mouse IL-6 ELISA Kit | R&D Systems | M6000B | Cytokine Quantification |
| Proteome Profiler Mouse Cytokine Array Kit | R&D Systems | ARY006 | Cytokine Quantification |
| Cyanine5 NHS ester | Lumiprobe | 23020 | In Vivo Biodistribution |
| Zeba™ Spin Desalting Columns | Thermo Scientific | A57762 | In Vivo Biodistribution |
| FSC22 Frozen Section Comp (9x118ml) Clear | Lecia | 3801480 | Tissue Cryosection |
| SuperFrost Plus™ Adhesion slides | Epredia™ | j1800amnz | Tissue Cryosection |
| Bovine Serum Albumin | TOCRIS | 5217 | Confocal Imaging of Liver Cryosection |
| Fluoromount-G | SouthernBiotech | 0100-01 | Confocal Imaging of Liver Cryosection |
| Cover glasses | Marienfeld | 0101242 | Confocal Imaging of Liver Cryosection |
| CD45 MicroBeads, mouse | Miltenyi Biotec | 130-052-301 | scRNA-seq |
| CellTracker™ Deep Red Dye | Invitrogen | C34565 | Apoptotic Cell Labeling |
| Lipopolysaccharides from *Escherichia coli* O111:B4 | Sigma-Aldrich | L4391-1mg | In Vitro Cell Treat |
| RPMI 1640 Medium | WELGENE | LM011-01 | BMDM Culture |
| Mouse M-CSF Recombinant Protein, PeproTech® | Gibco | 315-02 | BMDM Culture |
| NSC 74859 | R&D Systems | 4655/50 | STAT3 inhibitor |
| Annexin V | AdipoGen Life Sciences | AG-40B-0005TS-T100 | Ex Vivo Macrophage Survival |
| Annexin Binding Buffer (5X) | Invitrogen | V13246 | Ex Vivo Macrophage Survival |
| pHrodo™ Red S. aureus BioParticles™ Conjugate for Phagocytosis | Invitrogen | P35361 | Macrophage-Mediated Peri-Portal Zonation |

Supplement Table 2. Antibody List

| Molecule | Conjugate | Vender | Catalog # | Concentration | Concentration used | Usage |
| --- | --- | --- | --- | --- | --- | --- |
| CD45.2 | PerCP/Cy5.5 | BioLegend | 109828 | 0.2mg/ml | 2ng/ml | FACS |
| Ly6G | BV605 | BioLegend | 127639 | 0.2mg/ml | 2ng/ml | FACS |
| CD11b | Alexa Fluor 700 | BioLegend | 101222 | 0.5mg/ml | 5ng/ml | FACS |
| CD11b | FITC | BioLegend | 101206 | 0.5mg/ml | 5ng/ml | FACS |
| F4/80 | BV421 | BioLegend | 123132 | 0.2mg/ml | 2ng/ml | FACS |
| Tim-4 | PE/Cyanine7 | BioLegend | 130010 | 0.2mg/ml | 2ng/ml | FACS |
| CD64 | FITC | BioLegend | 139316 | 0.5mg/ml | 5ng/ml | FACS |
| CD31 | FITC | BioLegend | 102406 | 0.5mg/ml | 5ng/ml | FACS |
| MERTK | PE | BioLegend | 151506 | 0.2mg/ml | 2ng/ml | FACS |
| C3aR | BV786 | BD Biosciences | 753362 | 0.2mg/ml | 2ng/ml | FACS |
| C3aR | PE | BD Biosciences | 568276 | 0.2 mg/ml | 2ng/ml | FACS |
| CD9 | BV786 | BD Biosciences | 740886 | 0.2mg/ml | 2ng/ml | FACS |
| CD9 | PE | BioLegend | 124805 | 0.2mg/ml | 2ng/ml | FACS |
| Mouse BD Fc Block |  | BD Biosciences | 553142 | 0.5mg/ml | 10ug/ml | FACS |
| C3aR | PE | BD Biosciences | 568276 | 0.2 mg/ml | 5ng/ml | Confocal |
| CD9 | PE | BioLegend | 124805 | 0.2mg/ml | 5ng/ml | Confocal |
| E-Cadherin | BV421 | BioLegend | 147319 | 0.2mg/ml | 5ng/ml | Confocal |
| F4/80 | Alexa Fluor 647 | BD Biosciences | 565853 | 0.2mg/ml | 5ng/ml | Confocal |
| Myc |  | abcam | Ab9106 | 1 mg/mL | 1:1000 | Western Blot |
| HIF1a |  | R&D System | MAB1536 | 1mg/ml | 1:1000 | Western Blot |
| ALIX |  | Santa Cruz | Sc-53540 | 200 µg/ml | 1:100 | Western Blot |
| TSG101 |  | abcam | Ab83 | 0.5 - 1.8 mg/mL | 1:100 | Western Blot |
| Calnexin |  | abcam | Ab22595 | 0.4 - 1 mg/mL | 1:100 | Western Blot |
| Mouse IgG | Peroxidase | Sigma | A4416-1ML |  | 1:3000 | Western Blot |
| Rabbit IgG | Peroxidase | Sigma | A6154-1ML |  | 1:3000 | Western Blot |

Supplement Table 3. RT-PCR Primer List

| Target gene | Forward or Reverse | Sequence |
| --- | --- | --- |
| h_GAPDH | Forward | TGT GGT CAT GAG TCC TCC CA |
|  | Reverse | CGA GAT CCC TCC AAA ATC AA |
| h_VEGF | Forward | TGT GCC CAC TGA GGA GTC C |
|  | Reverse | GGT TTG ATC CGC ATA ATC TGC |
| h_GLUT1 | Forward | TGT GGG CCT TTT CGT TAA CC |
|  | Reverse | ATC ATC AGC ATT GAA TTC CGC |
| m_28s | Forward | CGA GAT TCC CAC TGT CCC TA |
|  | Reverse | GGG GCC TTC CCA CTT ATT CTA |
| m_Nr1h3 | Forward | AGG AGT GTC GAC TTC GCA AA |
|  | Reverse | CTC TTC TTG CCG CTT CAG TTT |
| m_C3ar1 | Forward | GGA AGC TGT GAT GTC CTG G |
|  | Reverse | CAC ACA TCT GTA CTC ATA TTG T |
| m_Cebpb | Forward | GCG CGA GCG CAA CAT C |
|  | Reverse | TGC TTG AAC AAG TTC CGC AG |
| m_Stat3 | Forward | AGG AGT CTA ACA ACG GCA GCC |
|  | Reverse | GTG GTA CAC CTC AGT CTC GAA G |
